# Hugin-AstA circuitry is a novel central energy sensor that directly regulates sweet sensation in *Drosophila* and mouse

**DOI:** 10.1101/2025.08.16.670687

**Authors:** Wusa Qin, Tingting Song, Zeliang Lai, Daihan Li, Liming Wang, Rui Huang

**Author notes:** These authors contribute equally to this work (Wusa Qin, Tingting Song).

## Abstract

Taste sensation plays a crucial role in shaping feeding behavior and is intricately influenced by internal states like hunger or satiety. Despite the identification of numerous neural substrates regulating feeding behavior, the central neural substrate that linked energy-sensing and taste sensation remained elusive. Here, we identified a novel neural circuitry that could directly sense internal energy state and modulate sweet sensation in the Drosophila brain. Specifically, a subset of neuropeptidergic neurons expressing hugin directly detected elevated levels of circulating glucose via glucose transporter Glut1 and ATP-sensitive potassium channel. Upon activation, these neurons released hugin peptide and activated downstream Allatostatin A (AstA)^+^ neurons via its cognate receptor PK2-R1. Subsequently, the activation of AstA^+^ neurons then directly inhibited sweet sensation via AstA peptide and its cognate receptor AstA-R1 expressed in sweet-sensing Gr5a^+^ neurons. We also showed that neuromedin U (NMU), the mammalian homolog of fly hugin, served as an energy sensor to suppress sweet sensation. Therefore, these data identify hugin^+^ neuron as a glucose-responsive central energy-sensing module that modulates sweet sensation across species.

## Introduction

Animal behavior is tightly coupled to internal metabolic states^1^. Hunger profoundly alters feeding-related behaviors, enhancing sensitivity to appetitive cues and promoting food-seeking, whereas satiety actively suppresses these responses to shift behavioral priorities^2,3^. Such state-dependent modulation allows animals to dynamically allocate resources to maintain energy homeostasis^4^. While the neural mechanisms that drive feeding during hunger have been extensively characterized^5^, the central circuits that sense satiety signals— specifically circulating glucose—to directly inhibit sensory processing remain less understood.

In both mammals and insects, “hunger signals” act as powerful accelerators for feeding. In the mammalian hypothalamus, AgRP neurons are activated by energy deficits to promote consumption^6,7^, while in *Drosophila*, dopaminergic and Neuropeptide F (NPF) pathways increase the gain of sweet-sensing gustatory neurons to drive sugar intake^8–10^. These systems explain how animals ramp up feeding motivation when energy is low. However, to prevent overconsumption, the brain must also possess precise “braking” mechanisms that detect elevated energy states and rapidly dampen sensory sensitivity.

Dietary sugar is a primary satiety signal^11^, yet the pathway linking central sugar sensing to behavioral inhibition remains elusive. In *Drosophila*, high-sugar diets are known to reduce sweet sensation via gut-derived signals or through central sensors like insulin-producing cells and DH44-expressing neurons that regulate general metabolic states^12–14^. Despite these advances, a critical gap remains: identifying a direct central sensor that detects acute rises in circulating glucose and acts specifically to inhibit the peripheral sweet-sensing machinery. Hugin-expressing neurons are promising candidates for this role, as they have been implicated in satiety-related feeding regulation^15,16^. Furthermore, Allatostatin A (AstA) has been proposed as a downstream signal capable of inhibiting sweet sensation^17^. However, whether and how these components form a unified circuit to sense glucose and apply the “brake” on feeding has not been established.

Here, we identify a complete neural circuitry that bridges this gap, directly linking internal glucose levels to the modulation of sweet taste sensation. We show that a specific subpopulation of hugin-expressing neurons functions as a central energy sensor, detecting elevated circulating glucose via the glucose transporter Glut1 and ATP-sensitive potassium channels. Upon activation, these neurons release hugin neuropeptide to activate downstream AstA-expressing neurons via the PK2-R1 receptor. Subsequently, AstA neurons directly inhibit sweet-sensing Gr5a^+^ gustatory neurons through AstA–AstA-R1 signaling. This pathway functions as a glucose-responsive inhibitory module whose endogenous activity is elevated in fed flies, thereby contributing to the suppression of sweet sensitivity after feeding. Finally, we demonstrate that Neuromedin U (NMU), the mammalian homolog of hugin^18^, similarly acts as a central energy sensor in mice to modulate sweet-responsive circuits, revealing a conserved neural strategy for coupling metabolic state to sensory processing across species.

## Results

### hugin^+^ neurons were a novel glucose sensor in the fly brain

Flies extend their proboscis in response to appetitive cues, a behavioral element known as the proboscis extension reflex (PER) ^20,51^. As previously observed, hunger significantly enhances this reflex^20^. In our study, we first replicated this phenomenon: In the state of hunger, flies exhibited elevated PER towards various concentrations of sugar compared to those under fed conditions (***Figure 1A-B***). These results confirmed the notion that sweet sensation was influenced by the internal energy state, elevated by starvation and suppressed by satiety ^20,52^. Sweet sensation in flies is mediated by sweet-sensing Gr5a^+^ neurons located on the proboscis^53^. Consistent with the behavioral effect, we also confirmed that Gr5a^+^ neurons exhibited elevated calcium responses to sugar under starvation conditions (***Figure supplementary 1A***).

**Figure 1.**
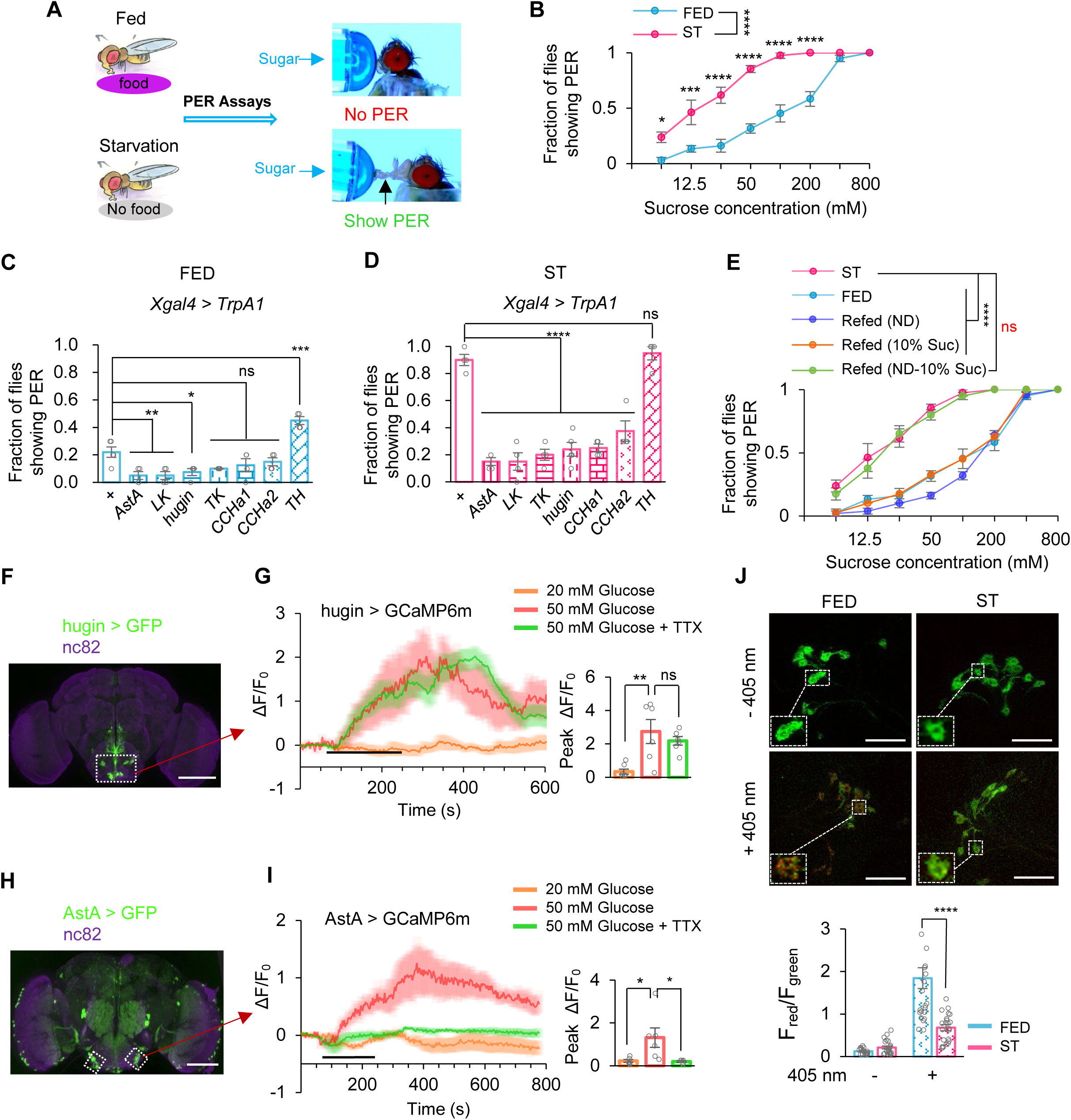
hugin^+^ neurons were a novel glucose sensor in the fly brain. (**A**) Experimental procedure schematic. Newly eclosed flies were gathered and provided with regular food for 5 days, followed by a 12-hour period of starvation. Proboscis Extension Response (PER) assays were conducted. (**B**) Fraction of flies showing PER to different concentrations of sucrose (two-way ANOVA; *,p=0.0349;***,p=0.002;****,p<0.0001; n=4 groups, each of 10 flies). (**C-D**) Fraction of flies of the indicated genotypes under 30 °C showing PER to 400 mM of sucrose (one-way ANOVA; *,p=0.0233;**,p=0.0054;***,p=0.001;****,p<0.0001; n=4-5 groups, each of 10 flies). (**E**) Fraction of indicated flies showing PER to different concentrations of sucrose (ND represent standard fly food) (two-way ANOVA; ****,p<0.0001; n=4-5 groups, each of 10 flies). (**F**) hugin expression in the brain, illustrated by mCD8::GFP expression driven by *hugin^GAL4^*. Scale bar represents 100 μm. (**G**) Representative traces and quantification of ex vivo calcium responses of hugin^+^ neurons during the perfusion of glucose with or without TTX (one-way ANOVA; **,p=0.002; n=6-7). Horizontal black bar represents the duration indicated glucose solution stimulation. (**H**) AstA expression in the brain, illustrated by mCD8::GFP expression driven by *AstA^GAL4^*. Scale bar represents 100 μm. (**I**) Representative traces and quantification of ex vivo calcium responses of AstA^+^ neurons during the perfusion of glucose with or without TTX (one-way ANOVA; *,p=0.0187 or 0.022; n=6). (**J**) Representative images of pre-photoconversion (pre-PC) and post-photoconversion (post-PC) CaMPARI signal in hugin-expressing neurons (upper). The Red:Green ratio represents intracellular Ca^2+^ concentrations (lower). Scale bar represents 100 μm (one-way ANOVA; ****,p<0.0001; n=18-23). ns, P > 0.05; *P < 0.05; **P < 0.01; ***P < 0.001; ****P < 0.0001. Student’s *t-test*, one-way and two-way ANOVA followed by post hoc test with Bonferroni correction were used for multiple comparisons when applicable.

Dopaminergic neurons in the subesophageal zone (SEZ) region of the fly brain acted as a hunger sensor, promoting sweet sensation via a specific dopamine receptor DopEcR expressed in Gr5a^+^ neurons ^20,52^. However, although the gene knockout of tyrosine hydroxylase (TH) ^54^, a key enzyme for dopamine biosynthesis, significantly suppressed sweet sensation in starved fruit flies (***Figure supplementary 1B***), these flies still showed a further reduction in sugar-induced PER under sated conditions (***Figure supplementary 1C-D***). These results suggest the existence of dopamine-independent mechanisms for sensing satiety and hunger in fruit flies that exerts modulatory effect on sweet sensation.

In the terminating phase of feeding behavior, satiety signals play a crucial role to cease food consumption and to ensure an appropriate amount of food ingestion. A number of neuropeptidergic neurons were identified as putative satiety signals ^36^. Notably, some of these neurons, including those secreting neuropeptides LK, TK, hugin, AstA, CCHa1, and CCHa2, are also distributed in the SEZ region ^55^. It has been reported that the sweet-sensing gustatory neurons primarily receive regulatory inputs from neurons in the SEZ region ^20,51^. Therefore, we hypothesized that these neuropeptidergic neurons might function as satiety signals, imposing inhibititory effect on sweet sensation after adequate food ingestion.

To test this hypothesis, we selectively expressed the temperature-sensitive ion channel TrpA1 in these neurons and activated them at 30 °C, triggering the release of corresponding neuropeptides. Indeed, activation of LK-, TK-, hugin-, AstA-, CCHa1-, and CCHa2-expressing neurons robustly suppressed sweet taste sensitivity in both fed and starved flies (***Figure 1C–D***; ***also see Figure supplementary 2***). Moreover, LK-, TK-, and hugin-expressing neurons exerted particularly strong inhibition under satiated conditions (***Figure 1C–D***; also see ***Figure supplementary 2***). These results reveal a multilayered, state-dependent neuropeptidergic network that finely tunes gustatory processing and feeding behavior. In contrast, activating dopaminergic neurons enhanced sweet perception in sated flies as expected (***Figure 1C***). In starved flies, activating dopaminergic neurons elicited no change in PER likely due to a ceiling effect (***Figure 1D***).

In mammals and flies, circulating sugar levels are directly linked to satiety/hunger state ^11,36,56^. We then asked whether changes in circulating sugar levels exerted an effect on sweet sensation. Activating IPCs in the fly brain released *Drosophila* Insulin-like Peptides (DILPs), the fly analog of mammalian insulin ^57^, into the hemolymph and reduced circulating sugar levels (***Figure supplementary 3A***). As a result, such manipulation enhanced sweet sensitivity of fed flies (***Figure supplementary 3B-C***). Therefore, it was plausible that changes in circulating sugar might be the driver of changes in sweet sensation.

Indeed, allowing starved flies to re-feed on normal fly diet or sucrose alone for 15 minutes both resulted in a significant decrease in sweet sensation (***Figure 1E***). Refeeding with multiple caloric sugars similarly suppressed sweet sensation (***Figure supplementary 4A***), indicating that energy intake contributes to the attenuation of sweet sensitivity. To examine whether sweet taste experience itself also plays a role, we refed starved flies with arabinose, a sweet but non-nutritive sugar. Arabinose feeding likewise led to a reduction in sweet sensitivity (***Figure supplementary 4B***), suggesting that sweet taste experience can contribute to the suppression of sweet perception, potentially through sensory adaptation. Consistent with this interpretation, responses to other sweet compounds, including trehalose and fructose, were also reduced following sucrose refeeding (***Figure supplementary 4C-D***). Conversely, when starved flies were supplied with fly diet deprived of sucrose (but with normal levels of polysaccharides, proteins, and lipids), there was no immediate change in sweet sensation after re-feeding (***Figure 1E***). These results demonstrate that sugar intake inhibits sweet sensation, probably via increasing circulating sugar levels.

Collectively, we thus hypothesized that the above neuropeptidergic neurons might directly sense an increase of circulating sugar levels and suppress sweet sensation in response. If so, these neurons should be activated by glucose, the main species of circulating sugars associated with feeding behavior in flies ^47^, at the concentrations resembling sated state (>50 mM in fed flies (***Figure supplementary 5***)). We thus conducted calcium imaging experiments in fly brains in an *ex vivo* preparation ^11^. Among all the six groups of neuropeptidergic neurons we examined in the SEZ, only hugin^+^ neurons and AstA^+^ neurons could be activated by 50 mM glucose (***Figure 1F-I, also see Figure supplementary 6***). Remarkably, each of these populations subdivided into anatomically distinct SEZ subclusters (***Figure supplementary 7A; Figure supplementary 8A***), yet glucose evoked Ca² responses occurred exclusively within a single subcluster (***Figure 1H,I; Figure supplementary 7B; Figure supplementary 8B***). This selective activation highlighted profound functional heterogeneity in nutrient sensing among these neuron groups. When direct synaptic transmission was blocked with TTX, hugin^+^ neurons, but not AstA^+^ neurons, still exhibited calcium responses towards 50 mM of glucose (***Figure 1G, green***), suggesting that hugin^+^ neurons are a direct glucose sensor. Furthermore, the level of circulating glucose dropped to ∼ 20 mM in starved flies (***Figure supplementary 5***). We found that 20 mM of glucose did not activate hugin^+^ and AstA^+^neurons (***Figure 1G and I, orange***), suggesting that these neurons can only be activated under sated conditions.

We also assessed the activity of hugin^+^ neurons *in vivo* using calcium-modulated photoactivatable ratiometric integrator (CaMPARI), a method employing a photoconvertible fluorescent protein that allows the imaging of integrated calcium activity ^58^, under both sated and starved conditions. In the CaMPARI assay, when neurons are activated, leading to an increase in free calcium ions, exposure to 405-nm light induces a permanent green-to-red photoconversion ^58^. We found an increased ratio of red/green fluorescence in hugin^+^ neurons in the sated state compared to that upon starvation (***Figure 1J***). These findings strongly imply that hugin^+^ neurons function as a glucose-sensitive energy sensor whose activity tracks internal metabolic state.

### Glucose activated hugin^+^ neurons via an ATP-sensitive potassium channel

We then investigated the cellular mechanism through which glucose could directly activate hugin^+^ neurons. In both mammals and flies, extracellular glucose is usually transported into the cytosol to activate glucose-sensitive neurons ^13,14,59^. When fly brains were pre-treated with phlorizin, a blocker of glucose transport ^13,14^, hugin^+^ neurons showed diminished calcium responses to glucose, suggesting activation of hugin^+^ neurons by glucose requires intracellular glucose (***Figure 2A***).

**Figure 2.**
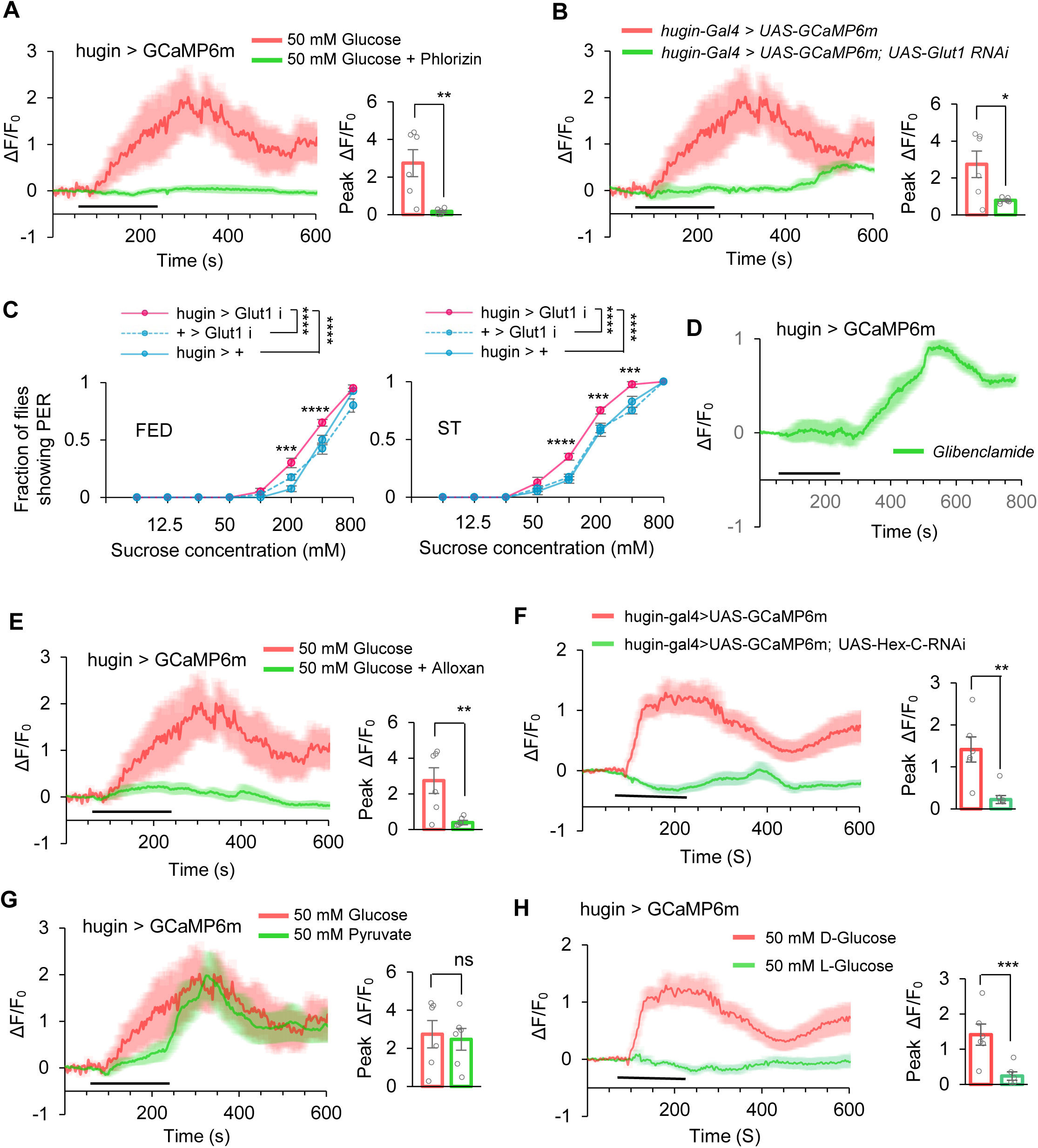
Glucose activated hugin^+^ neurons via Gltu1 and K_ATP_ channel. (**A**) Representative traces and quantification of ex vivo calcium responses of hugin^+^ neurons during the perfusion of glucose with or without phlorizin (1mM),(t-test;**,p=0.0052; n=6-7). Horizontal black bar represents the duration indicated glucose solution stimulation. (**B**) Representative traces and quantification of ex vivo calcium responses of hugin^+^ neurons in indicated flies during the perfusion of glucose (t-test;*,p=0.022; n=6). (**C**) Fraction of flies of the indicated genotypes showing PER to different concentrations of sucrose (two-way ANOVA; ****,p<0.0001;n=4 groups, each of 10 flies). (**D-E**) Representative traces and quantification of ex vivo calcium responses of hugin^+^ neurons during the perfusion of glibenclamide (100 μM, D), glucose with alloxan (10 μM, E) (t-test;**,p=0.0091; n=6). (**F**) Representative traces and quantification of ex vivo calcium responses of hugin^+^ neurons in indicated flies during the perfusion of glucose (t-test;**,p=0.0011; n=6). (**G**) Representative traces and quantification of ex vivo calcium responses of hugin^+^ neurons during the perfusion of pyruvate (50 mM), (t-test;ns,p>0.05; n=6). (**H**) Representative traces and quantification of ex vivo calcium responses of hugin^+^ neurons during the perfusion of D-glucose and L-glucose (50 mM), (t-test;***,p=0.0045; n=6) ns, P > 0.05; *P < 0.05; **P < 0.01; ****P < 0.0001. Student’s t-test and two-way ANOVA followed by post hoc test with Bonferroni correction were used for multiple comparisons when applicable.

The transport of glucose mediated by glucose transporter 1 (Glut1) can activate certain neurons in the fruit fly brain ^13,14^. By specifically knocking down Glut1 expression in hugin^+^ neurons, we observed a significant reduction in calcium responses to glucose (***Figure 2B***). Consistently, these flies showed a slight yet significant increase in their PER responses to glucose (***Figure 2C***). Again, these results further confirm that glucose needs to be transported into hugin^+^ neurons to modulate neuronal activity and behavioral output.

In pancreatic β-cells and certain fly neurons ^13,59,60^, intracellular glucose modulates cell excitability primarily through its effect on intracellular energy metabolism and ATP-sensitive potassium channels (K_ATP_). We pre-treated fly brains with a K_ATP_ inhibitor (glibenclamide) and observed a significant calcium response in hugin^+^ neurons (***Figure 2D***). Hexokinase, a key enzyme in glucose metabolism that generates ATP, also plays a crucial role in this process. When hexokinase activity was restricted by alloxan, glucose-induced calcium response in hugin^+^ neurons was significantly suppressed (***Figure 2E***). Similarly, when hexokinase was specifically knocked down in hugin^+^ neurons, a comparable reduction in glucose-induced calcium responses was observed (***Figure 2F***). Pyruvate, the end product from glycolysis, can lead to ATP generation via the tricarboxylic acid cycle. We also found that pyruvate stimulation led to a significant calcium response in hugin^+^ neurons (***Figure 2G***). As a specificity control, nonmetabolizable L-glucose failed to activate hugin neurons (***Figure 2H***).

Collectively, these findings suggest that extracellular glucose directly enters hugin^+^ neurons through specific glucose transporter and subsequently modulate the activity of K_ATP_ through ATP production, ultimately leading to neuronal depolarization. We note that blocking glucose transport or metabolism could, in principle, nonspecifically affect neuronal function, but the rapid and selective nature of the observed responses argues against a generalized loss of excitability.

### AstA^+^ neurons were direct downstream target of hugin^+^ neurons

Since hugin^+^ neurons were directly involved in glucose sensing and regulated sweet sensation (***Figure 1G and Figure3A-B***), we sought to identify their downstream target. We then asked whether the hugin peptide had a direct effect on sweet sensation. Upon microinjection of synthetic hugin peptide, we observed a notable reduction in sweet sensation under both sated and starved states (***Figure 3C***). In line with this observation, following the injection of hugin peptide, we observed a significant reduction of sugar-induced calcium responses in sweet-sensing Gr5a^+^ neurons (***Figure 3D***). Thus, the inhibitory function of hugin^+^ neurons on sweet sensation was likely mediated by the secretion of hugin peptide.

**Figure 3.**
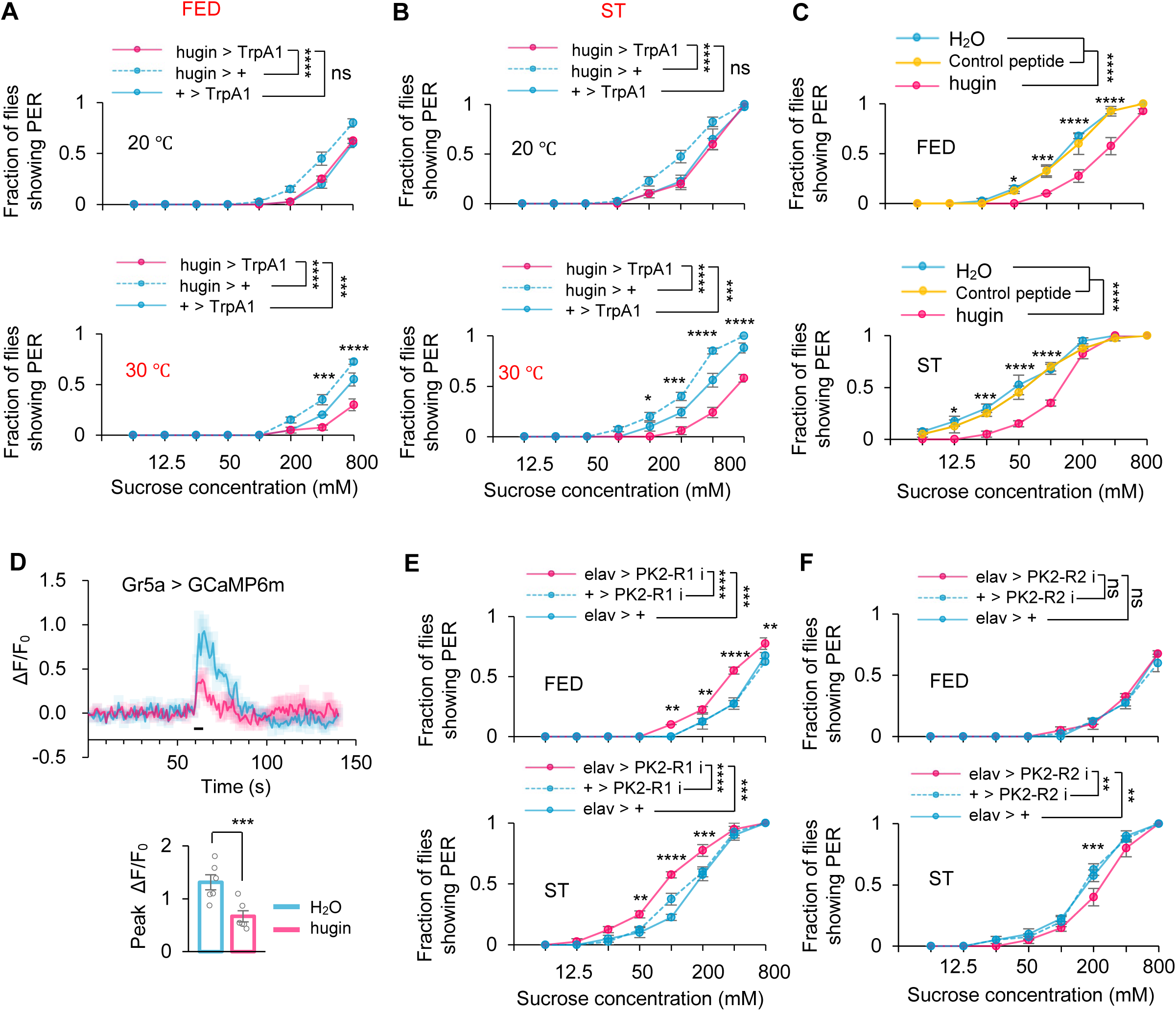
hugin^+^ neurons suppressed sweet taste through PK2-R1. (**A-B**) Fraction of flies of the indicated genotypes and environmental temperatures showing PER to different concentrations of sucrose (two-way ANOVA; *,p=0.0153 or 0.0237; ****,p<0.0001;n=4-5 groups, each of 10 flies). (**C**) Fraction of flies showing PER to different concentrations of sucrose (two-way ANOVA; *,p=0.0486 or 0.0438; ***,p=0.0007 or 0.0001; ****,p<0.0001; n=4 groups, each of 10 flies). Flies were injected with saline or synthetic hugin for 30 minutes before the assay. (**D**) Representative traces (upper) and quantification (lower) of peak calcium transients of Gr5a^+^ neurons in indicated flies upon 5% sucrose after injection of synthetic hugin (t-test;***,p=0.0047; n=6). Horizontal black bar represents the duration sucrose stimulation. (**E-F**) Fraction of flies of the indicated genotypes showing PER to different concentrations of sucrose (two-way ANOVA; **,p=0.0055 or 0.0016 or 0.003; ***,p=0.0002 or 0.0005; ****,p<0.0001;n=4 groups, each of 10 flies)ns, P > 0.05; *P < 0.05; ***P < 0.001; ****P < 0.0001. Two-way ANOVA followed by post hoc test with Bonferroni correction was used for multiple comparisons when applicable.

There were two hugin receptors, PK2-R1 and PK2-R2, in fruit flies ^61^. We knocked down their expression by RNAi in the nervous system and then tested flies’ PER to sugar. Knocking down PK2-R1 but not PK2-R2 significantly enhanced sweet perception in both sated and starved states (***Figure 3E-F***), suggesting a crucial role of PK2-R1 in receiving the input of hugin^+^ neurons in satiety sensing. To validate this finding, we further tested PK2-R1 null mutants and observed a similar enhancement in sweet sensitivity (***Figure supplementary 9***). Although PK2-R2 knockdown did not enhance PER, we observed a modest but reproducible reduction in PER in starved flies (***Figure 3F***). This suggests potential functional heterogeneity within hugin signaling pathways. Given the anatomical diversity of hugin neurons, distinct subpopulations may engage different receptor-dependent mechanisms that exert complementary or even opposing effects on feeding behavior. Further dissection of PK2-R2-dependent signaling will be required to clarify this complexity.

Interestingly, we did not observe any expression of PK2-R1 in the proboscis where gustatory sensory neurons were distributed, indicating that sweet-sensing Gr5a^+^ neurons could not directly receive input from hugin^+^ neurons via PK2-R1 (***Figure supplementary 10***).AstA^+^ neurons served as a plausible relay between hugin^+^ neurons and sweet-sensing Gr5a^+^ neurons (***Figure 4A-B***), especially based on a clear co-expression of hugin receptor PK2-R1 and AstA in the SEZ region of the fly brain (***Figure 4C***). Notably, these double-labeled neurons corresponded precisely to the AstA subcluster that exhibited glucose specific Ca² responses (***Figure 1H***). We also used optogenetic tools to further confirm that hugin^+^ neurons acted upstream of AstA^+^ neurons. We expressed the light-sensitive neuronal activator CsChrimson in hugin^+^ neurons and calcium indicator GCaMP6m in AstA^+^ neurons, respectively ^62^. As shown in Figure 4D, opto-activation of hugin^+^ neurons led to a robust calcium transient in AstA^+^ neurons.

**Figure 4.**
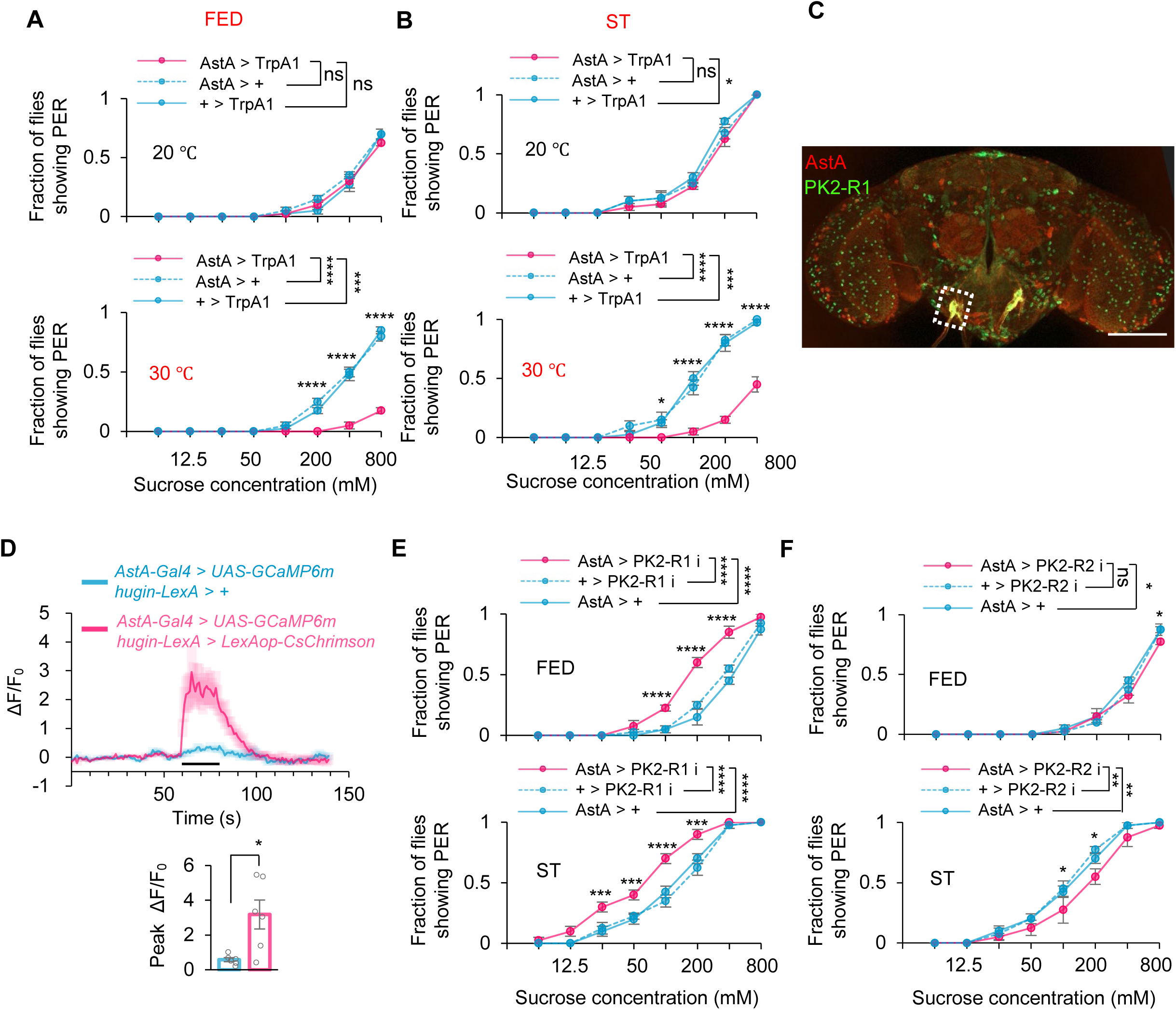
hugin^+^ neurons activated AstA^+^ neurons through PK2-R1. (**A-B**) Fraction of flies of the indicated genotypes and environmental temperatures showing PER to different concentrations of sucrose (two-way ANOVA; *,p=0.043; ***,p=0.0007 or 0.0001; ****,p<0.0001;n=4-5 groups, each of 10 flies). (**C**) Co-localization (dashed box) of PK2-R1^+^ neurons (green) and AstA^+^ neurons (red) in the SEZ region. Scale bar represents 100 μm. (**D**) Representative traces (upper) and quantification (lower) of peak calcium transients of AstA^+^ neurons after the photo-activation of hugin^+^ neurons (t-test;*,p=0.0114; n=6) from in vivo calcium imaging. Horizontal black bar represents the duration of red-light stimulation. (**E-F**) Fraction of flies of the indicated genotypes showing PER to different concentrations of sucrose (two-way ANOVA; *,p=0.0486 or 0.033 or 0.0224;**,p=0.0043 or 0.0011; ***,p=0.0001 or 0.0008; ****,p<0.0001; n=4 groups, each of 10 flies). ns, P > 0.05; *P < 0.05; **P < 0.01; ***P < 0.001; ****P < 0.0001. Student’s *t-test* and two-way ANOVA followed by post hoc test with Bonferroni correction were used for multiple comparisons when applicable.

We then investigated whether the hugin-AstA circuitry affected sweet sensation. As expected, knocking-down PK2-R1 (but not PK2-R2) in AstA^+^ neurons led to a significant increase in sugar-induced PER (***Figure 4E-F***). Therefore, AstA^+^ neurons are the direct downstream target of hugin^+^ neurons in satiety sensing. hugin^+^ neurons can directly sense elevated circulating glucose levels, promote the release of neuropeptide hugin, and then activate AstA^+^ neurons via its cognate receptor PK2-R1.

hugin^+^ neurons are distributed in both the brain and the ventral nerve cord (VNC). The hugin^+^ neurons in the VNC sense sugar and inhibit feeding behavior by suppressing the activity of DH44 neurons ^48^. To determine which hugin^+^ neurons regulate the activity of AstA neurons and sweet sensation, we performed the following experiments: First, we physically severed the neural connection between the brain and the VNC. Under this condition, activation of Hugin neurons still significantly reduced PER responses ***(Figure supplementary 11A–B)***, indicating that hugin neurons within the brain are sufficient to modulate sweet perception. Second, in optogenetic experiments using isolated brain preparations, light stimulation of hugin neurons elicited robust calcium responses in AstA neurons ***(Figure supplementary 11C)***, demonstrating a direct functional coupling between brain Hugin neurons and AstA neurons.

Notably, comparison of intact and severed preparations revealed that the magnitude of AstA neuronal activation was greater when VNC neurons were preserved, suggesting that VNC hugin neurons may provide additional modulatory input that enhances AstA activation. Together, these results indicate that Hugin neurons in the SEZ constitute a primary pathway linking sugar sensing to AstA activation and PER regulation, while VNC hugin neurons contribute to the overall strength of hugin signaling.

### AstA^+^ neurons directly inhibited Gr5a^+^ neurons

Activation of AstA^+^ neurons suppressed sweet sensation, resembling the effect of activating hugin^+^ neurons ^17^(***Figur 4A-B***). We also evaluated the behavioral effect of AstA peptide. Similar to hugin, the microinjection of AstA resulted in a reduction in sweet sensitivity in fruit flies (***Figure 5A***) and the activity of Gr5a^+^ neurons (***Figure 5B***). We then speculated whether AstA^+^ neurons and AstA peptide might function directly on Gr5a^+^ neurons. The cell bodies of sweet-sensing Gr5a^+^ neurons are located in the proboscis, whereas their axons extend to the SEZ region of the fly brain ^63^. We observed distributions of the projections of AstA^+^ neurons and Gr5a^+^ neurons both in the SEZ region (***Figure 5C***), suggesting that Gr5a^+^ neurons might directly receive AstA signal in this region.

**Figure 5.**
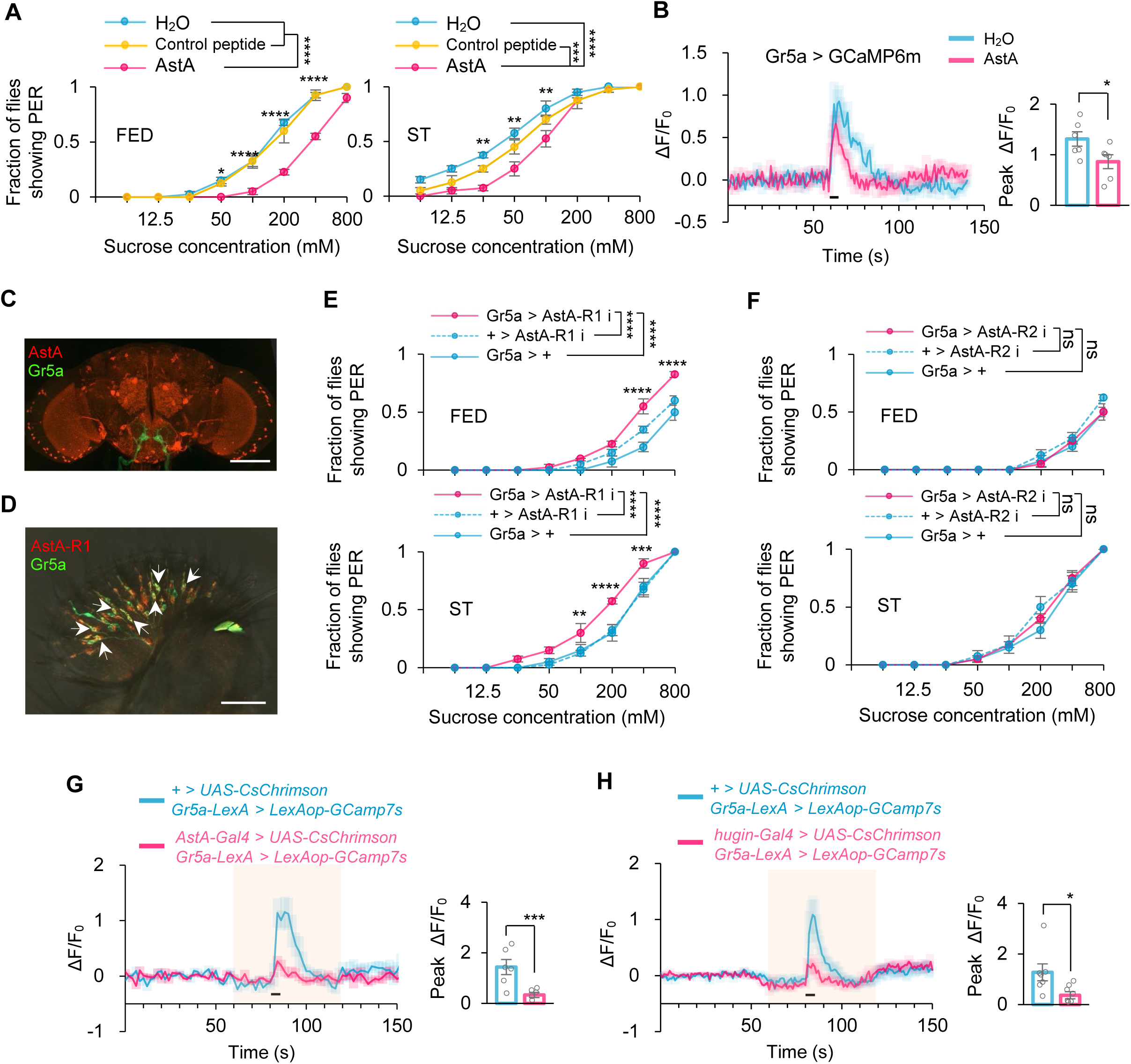
AstA^+^ neurons inhibited Gr5a^+^ neurons through AstA-R1. (A) Fraction of flies showing PER to different concentrations of sucrose (two-way ANOVA; *,p=0.024;**,p=0.0072 or 0.002; ***,p=0.0002; ****,p<0.0001; n=4 groups, each of 10 flies). Flies were injected with saline or synthetic AstA for 30 minutes before the assay. (B) Representative traces (left) and quantification (right) of peak calcium transients of Gr5a^+^ neurons in indicated flies upon 5% sucrose after injection of synthetic AstA (t-test;*,p=0.0476; n=6). Horizontal black bar represents the duration sucrose stimulation. (**C**) Localization of Gr5a^+^ neurons (green) and AstA^+^ neurons (red) in the brain. Scale bar represents 100 μm. (**D**) Co-localization of Gr5a^+^ neurons (green) and AstA-R1^+^ neurons (red). Arrows point to the neurons co-expressing the two receptors. Scale bar represents 50 μm. (**E-F**) Fraction of flies of the indicated genotypes showing PER to different concentrations of sucrose (two-way ANOVA **,p=0.0088; ***,p=0.0004; ****,p<0.0001; n=4 groups, each of 10 flies). (**G-H**) Representative traces (left) and quantification (right) of peak calcium transients of Gr5a^+^ neurons in indicated flies upon 5% sucrose during the process of photo-activation of AstA^+^ neurons (G, n=6; t-test;**,p=0.0058) and hugin^+^ neurons (H, n=7-8;t-test;*,p=0.0351). Shadows represents the duration of red-light stimulation. Horizontal black bar represents the duration of sucrose stimulation. ns, P > 0.05; *P < 0.05; ***P < 0.001; ****P < 0.0001. Student’s *t-test* and two-way ANOVA followed by post hoc test with Bonferroni correction were used for multiple comparisons when applicable.

AstA has two cognate receptors in fruit flies, AstA-R1 and AstA-R2 ^64^. Co-labeling experiments in the proboscis confirmed the expression of AstA-R1 in Gr5a^+^ neurons (***Figure 5D, arrows***), while AstA-R2 was not expressed in the proboscis (***Figure supplementary 12***). Functionally, knockdown of AstA-R1 specifically in Gr5a^+^ neurons, but not that of AstA-R2, significantly enhanced sweet sensation of flies (***Figure 5E-F***). Similar results were observed in AstA-R1 knockout flies (***Figure supplementary 13***).

Using optogenetic tools, we also confirmed the function of hugin^+^ neurons and AstA^+^ neurons on Gr5a^+^ neurons. Activating hugin^+^ neurons and AstA^+^ neurons both significantly reduced sugar-induced responses of Gr5a^+^ neurons (***Figure 5G-H***). Collectively, these results suggest that satiety-sensing hugin-AstA circuitry can directly modulate sweet sensation of flies via suppressing the activity of Gr5a^+^ gustatory neurons.

### hugin–AstA circuitry suppressed food consumption

Collectively, the hugin-AstA circuitry senses satiety signals and inhibits sweet sensation. Considering the crucial role of sweet sensation in regulating feeding behavior, we hypothesized that hugin-AstA circuitry was also involved in the regulation of food consumption.

We expressed a temperature-sensitive blocker of synaptic transmission (Shibire^ts1^) in hugin^+^ neurons or AstA^+^ neurons ^65^ and assayed the feeding behavior of flies under both permissive temperature (***Figure6A, blue***) and non-permissive temperature (***Figur 6A, red***). In contrast to the suppression of sweet sensitivity observed upon thermogenetic activation of these neurons, silencing hugin neurons led to enhanced sweet-driven feeding behavior (***Figure 6B***). Consistently, inhibiting synaptic transmission in either hugin or AstA neurons resulted in a significant increase in food consumption at the non-permissive temperature compared to controls (***Figure 6C–D***).

**Figure 6.**
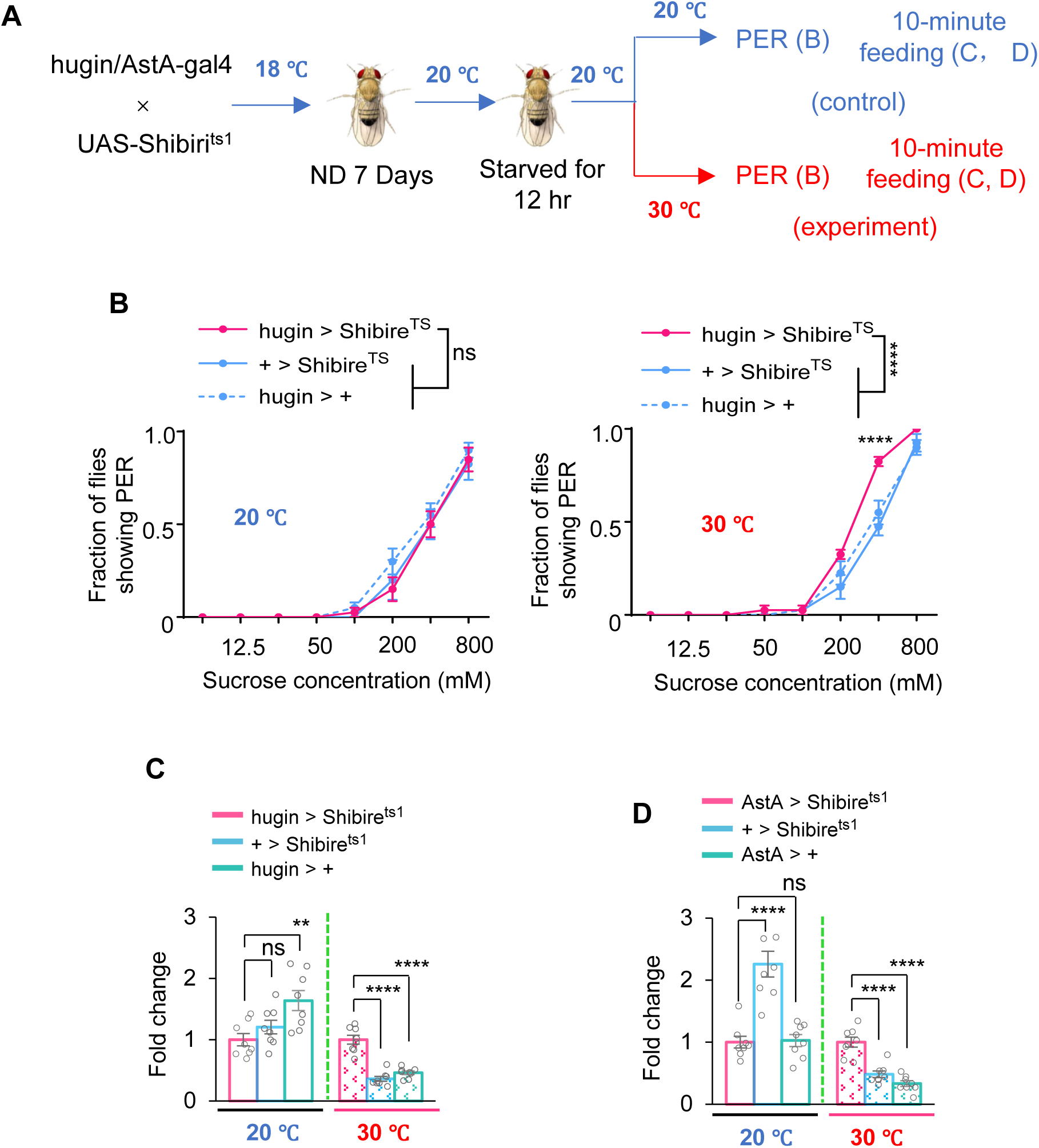
hugin-AstA circuitry inhibited feeding behavior. (A) The illustration of experimental design in this Figure. Note that all flies were maintained at 20°C before the feeding assay to prevent neuronal silencing by Shibire^ts1^ during these procedures. (B) Fraction of flies of the indicated genotypes and environmental temperatures showing PER to different concentrations of sucrose (two-way ANOVA; ****,p<0.0001; n=4 groups, each of 10 flies). (**C-D**) Food consumption of flies of the indicated genotypes and environmental temperatures (one-way ANOVA; **,p=0.004; ****,p<0.0001; n=8 groups, each of 4 flies). Food consumption is presented as fold change relative to the corresponding genetic control.

We also examined the role of hugin and AstA receptors in feeding behavior by testing *PK2-R1* and *AstA-R1* gene knockout flies. The results showed that knockout of both receptors enhanced feeding behavior (***Figure supplementary 14A***), consistent with increased sweet perception upon genetic manipulation of these two receptors (***Figure 4-5***). Moreover, these receptor knockout flies exhibited increased lipid storage, which was likely a consequence of their enhanced feeding behavior (***Figure supplementary 14B***). Additionally, we knocked down the expression of AstA-R1 in sweet sensing Gr5a^+^ neurons, and found that these flies exhibited a significant increase in food consumption (***Figure supplementary 14C***), also likely due to the enhanced sweet sensation (***Figure 5E***).

### Mammalian homolog of hugin was also a putative satiety sensor

Our present work identified hugin-AstA circuitry as a novel satiety sensor and feeding suppressor in fruit flies. Fruit flies and mammals share highly conserved regulatory elements on feeding behavior and energy homeostasis ^51,66,67^. We thus asked whether this novel satiety-sensing mechanism was also conserved in the mammalian system,

In rodent models, neuromedin U (NMU), a analogues of hugin, was also found to be an appetite suppressor ^15,18^. We found that synthetic NMU could suppress sweet perception when injected into flies’ thorax (***Figure supplementary 15***), highlighting the analogy of fly hugin and mammalian NMU. In mouse models, we observed a significant increase in NMU levels in the blood following glucose feeding (***Figure 7A***). Using the sucrose preference test (SPT) to measure sweet preference, we noted that NMU injection reduced sugar preference in mice, phenocopying the effect of glucose feeding (***Figure 7B-C***). Conversely, NMU knockout enhanced sweet sensation in mice (***Figure 7D***). These data demonstrated that both NMU gene and protein plays a crucial role in regulation of sweet sensation.

**Figure 7.**
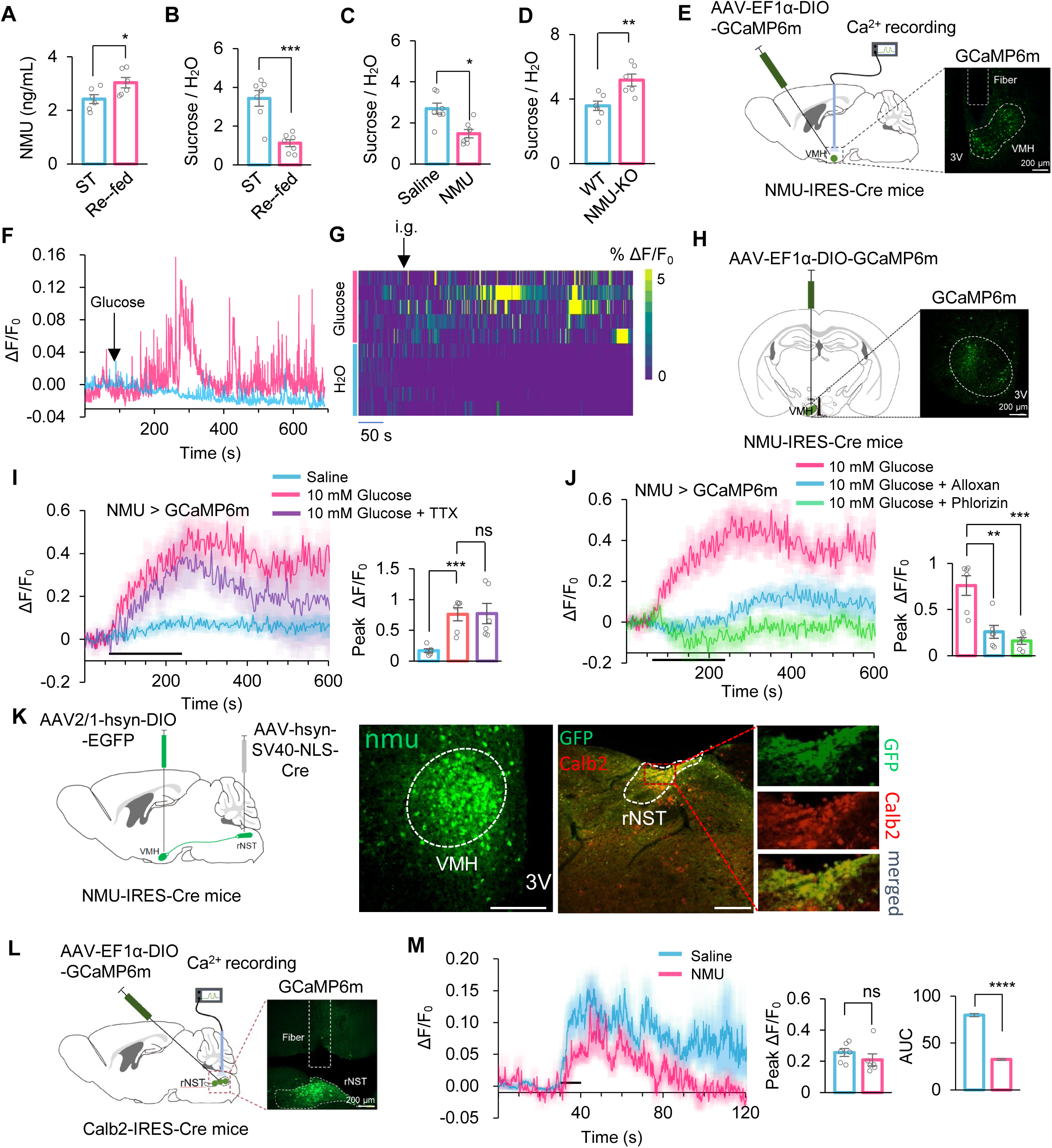
NMU^+^ neurons were the central energy sensor in mouse. **(A)** Blood NMU levels of starved mice re-fed with 20% sucrose (t-test; *, p=0.0373; n=6). **(B)** Two-bottle preference tests for starved mice re-fed with sucrose (t-test; ***, p=0.0002; n=7). (**C**) Two-bottle preference tests for starved mice with or without intraperitoneal injection of NMU peptide (YFLFRPRN-NH_2_, 4.5μm/kg) (t-test; *, p=0.0293; n=7). (**D**) Two-bottle preference tests for indicated starved mice (t-test; **, p=0.0073; n=6). (**E**) Fiber photometry to record the calcium dynamics of NMU^+^ neurons in the VMH with GCaMP6m (left) and representative IHC image of expressing GCaMP6min VMH (right). (**F-G**) Representative trace (left) and heatmaps (right, n=5) showing calcium dynamics of NMU^+^ neurons in the VMH in response to gastric glucose infusion (20% sucrose, 200 μl), which elevates circulating glucose levels independently of oral sensory stimulation.. (**H**) Experimental approach to assess calcium signaling in NMU^+^ neurons in the VMH in vitro (left) and representative IHC image of expressing GCaMP6min VMH (right). (**I-J**) Representative traces and quantification of ex vivo calcium responses of NMU^+^ neurons during the perfusion of glucose with or without TTX (I), Alloxan (J) and Phlorizin (J) (one-way ANOVA; **, p=0.0049;***, p=0.0001 or 0.0006; n=6-7). Horizontal black bar represents the duration indicated glucose solution stimulation. (**K**) Anterograde trans-synaptic tracing of downstream targets of NMU neurons. NMU neurons were labeled by GFP expression following injection of a Cre-dependent AAV2/1-DIO-GFP into NMU-Cre mice. This virus undergoes anterograde trans-synaptic transfer to postsynaptic neurons. To enable GFP expression specifically in downstream target regions, an AAV-Cre virus was locally injected into the rNST. As a result, postsynaptic neurons in the rNST receiving input from NMU neurons were labeled by GFP delivered anterogradely from upstream NMU neurons. Representative images show that GFP-labeled downstream neurons in the rNST colocalize with Calb2 immunoreactivity, indicating that NMU neurons preferentially target Calb2 rNST neurons. (**L**) Fiber photometry to record the calcium dynamics of Calb2^+^ neurons in the rNST with GCaMP6m (left) and representative IHC image of expressing GCaMP6m in rNST (right). (**M**) Representative traces and quantification of calcium responses of Calb2^+^ neurons during glucose licking under physiological feeding conditions (500 mM sucrose), with or without NMU administration, assessing downstream modulation of sweet-responsive brainstem circuits(t-test; ****, p<0.0001; n=6). ns, P > 0.05; *P < 0.05; **P < 0.01; ***P < 0.001; ****P < 0.0001. Student’s *t-test* and one-way ANOVA followed by post hoc test with Bonferroni correction was used for multiple comparisons when applicable.

Neuropeptide NMU is secreted by a group of neurons located in the VMH region of hypothalamus ^18^. To investigate whether NMU^+^ neurons functioned as an energy sensor in the mouse brain, we injected AAV-EF1α-DIO-GCaMP6m-WPRE into the VMH of NMU-Cre mice, specifically expressing GCaMP6m in NMU^+^ neurons. We then monitored the dynamic activity of VMH ^NMU^ neurons during glucose stimulation by using fiber photometry (***Figure 7E***). As expected, these neurons showed significant calcium responses following glucose administration in mice (***Figure 7F-G***). To confirm the specificity of GCaMP6m expression in NMU neurons, we co-expressed a Cre-dependent mCherry reporter in the VMH of NMU-Cre mice and performed fluorescence in situ hybridization for NMU mRNA. Colocalization analysis demonstrated that GCaMP6m fluorescence was restricted exclusively to NMU cells (***Figure supplementary 16A***).

Next, we conducted calcium imaging of brain slices *in vitro* to further assess these neurons’ ability to sense glucose (***Figure 7H***). Similar to hugin neurons in the fly brain, VMH^NMU^ neurons in the mouse brain exhibited strong calcium responses to glucose stimulation, which could not be inhibited by TTX (***Figure 7I***), indicating their direct glucose-sensing capability. Also similar to flies, glucose activated these mouse neurons via entering the cells and modulating K_ATP_ channels, as evidenced by the inhibition of calcium responses by alloxan and phlorizin (***Figure 7J***). While supraphysiological glucose concentrations were used in slice recordings to ensure reliable neuronal activation under ex vivo conditions, complementary in vivo recordings demonstrate that VMH NMU neurons are engaged by physiological sugar ingestion.

Calbindin 2-positive neurons (Calb2) in the rostral nucleus of the solitary tract (rNST) selectively respond to sweet tastants, facilitating sweet signal from the periphery into the brain ^68^. Because attenuated sweet sensation should correspondingly diminish central sweet signal transmission, we asked whether NMU modulates this relay. To assess potential anatomical connectivity between VMH NMU neurons and rNST Calb2 neurons, we employed a dual-virus tracing strategy. In NMU-Cre mice, we injected AAV9-EF1α-DIO-EGFP into the VMH, which revealed dense GFP projections in close apposition to Calb2 neurons in the rNST (***Figure supplementary 16B***). To further confirm this projection is monosynaptic, we combined anterograde tracing by co-injecting AAV2/1-EF1α-DIO-EGFP into the VMH and AAV2/1-hsyn-SV40-NLS-Cre into the rNST. This approach yielded GFP expression in rNST Calb2 neurons, supporting a direct VMH→rNST NMU →Calb2 synaptic link (***Figure 7K***).

To evaluate the functional consequence of NMU on Calb2 neuronal activity during sweet stimulation, we delivered AAVs encoding Cre-dependent GCaMP6m into the rNST of Calb2-Cre mice, thereby enabling real-time calcium imaging in Calb2 neurons (***Figure supplementary 16C***). Using fiber photometry, we recorded calcium responses to sugar stimuli, both with and without NMU injection (***Figure 7L***). We found that NMU injection elicited a significant, albeit partial, suppression of calcium responses to sugar ***(Figure 7M)***. Although the magnitude of this reduction was modest compared to the complete silencing of activity, the effect was statistically robust and aligned with the behavioral phenotypes, indicating that NMU acts to dampen sweet taste perception during feeding.

These findings indicate that NMU^+^ neurons can indeed be a putative satiety sensor in the mouse brain that suppresses sweet perception, shedding light on understanding the conserved neural mechanism of feeding regulation in mammals including human.

## Discussion

In this study, we identify a neural pathway that directly couples internal glucose levels to the regulation of sweet taste perception and feeding behavior. In *Drosophila*, circulating glucose activates a specific subset of hugin neurons, which signal to AstA neurons to suppress the activity of peripheral sweet-sensing Gr5a neurons, thereby reducing sweet sensitivity and terminating feeding. This pathway establishes a direct neural link between internal energy state and early sensory processing (***Figure supplementary 17***).

A notable feature of this circuit is that its manipulation produces significant but relatively modest changes in sweet sensitivity compared with the dramatic enhancement observed under starvation^52,69^. This difference reflects the distinction between a specific satiety mechanism and the global physiological state of hunger. Starvation is a systemic crisis that engages multiple hunger-promoting pathways, including dopaminergic and NPF signaling, which broadly elevate motivational drive and sensory gain^10,70^. In contrast, the hugin–AstA pathway functions as a glucose-dependent inhibitory branch whose endogenous activity is elevated in fed states but remains functionally capable of modulating sweet sensitivity across feeding conditions.. Disrupting this inhibitory pathway prevents full suppression of sweet sensitivity after feeding but does not recreate the potent hunger drive generated by dopaminergic activation. Thus, energy homeostasis arises from the coordinated action between hunger accelerators and satiety brakes.

This framework reconciles our findings with previous reports that silencing hugin neurons does not alter proboscis extension in starved flies^20^. Although endogenous hugin neuronal activity is reduced during starvation, our data show that further manipulation of this pathway can still bidirectionally influence feeding behavior under starved conditions. This indicates that the hugin–AstA circuitry is not exclusively restricted to fed states, but rather represents a state-modulated inhibitory system whose baseline tone shifts with circulating glucose levels. Under starvation, reduced endogenous activity may partially relieve this inhibitory influence; however, the circuit retains functional capacity to modulate sweet sensitivity when experimentally perturbed. In contrast, preventing the starvation-induced reduction of hugin activity—such as by thermogenetic activation—significantly suppresses hunger-enhanced sweet sensitivity. These results indicate that the dynamic downregulation of hugin neuron activity is a critical component of the state-dependent modulation of taste. Therefore, rather than functioning as a strictly “satiety-specific brake,” the hugin–AstA axis is better conceptualized as a glucose-responsive inhibitory module that biases sweet perception according to metabolic state. Its activity increases following feeding to dampen sensory gain, while decreased activity during starvation permits heightened sweet sensitivity. Such graded modulation enables flexible integration of internal energy status with sensory processing.

Our data further reveal functional heterogeneity within peptidergic populations. Although hugin and AstA neurons are broadly distributed in the central nervous system (including the subesophageal zone and ventral nerve cord), only specific subpopulations respond to elevated glucose^43,48^. While descending inputs from the VNC may potentiate signaling, our data suggest that the glucose-responsive hugin neurons in the SEZ constitute a dedicated energy-sensing module. This heterogeneity likely explains both the partial behavioral effects of global manipulations and the ability of hugin neurons to regulate diverse behaviors beyond feeding, such as locomotion.

Circulating sugars represent a complex internal signal. In *Drosophila*, trehalose serves as the major circulating sugar supporting stress resistance, whereas glucose functions as the primary, rapidly mobilized energy substrate^71^. Unlike receptor-based sensors (e.g., Gr43a for fructose)^72^, Hugin neurons rely on intracellular glucose metabolism and K^ATP^ channel activity to modulate neuronal excitability. This mechanism ensures that the circuit activity is tightly coupled to cellular energy status. Consistent with this, feeding with fructose, trehalose, or non-nutritive sweeteners only weakly suppressed sweet sensitivity within the acute post-feeding interval, whereas sucrose—which rapidly restores hemolymph glucose—robustly activated this pathway. Thus, our work identifies glucose as the key metabolic signal that engages this central brake.

The presence of multiple sugar-sensing systems in *Drosophila* reflects functional specialization rather than redundancy. Peripheral and gut-derived signals regulate feeding over longer timescales, while central hunger circuits adjust sensory gain during deprivation. The hugin–AstA circuit adds a distinct layer by acting as a rapid central brake that prevents overconsumption when energy stores are sufficient.

Strikingly, we find that this circuit logic is conserved in mammals. Although NMU has been implicated in feeding suppression^73,74^, whether NMU neurons sense internal nutrients directly had remained unclear. Our data demonstrate that VMH NMU neurons are activated by elevations in internal glucose and form a functional circuit with sweet-responsive Calb2 neurons in the rostral nucleus of the solitary tract (rNST). Gastric glucose infusion experiments established that NMU neurons respond to internal metabolic signals independently of oral sensory input, while physiological sugar ingestion confirmed that this pathway actively suppresses downstream sweet-driven activity. Together, these findings define a previously unrecognized NMU-centered circuit linking internal glucose sensing to the modulation of early taste processing. All mouse experiments in this study were performed in male animals; whether sex-specific differences exist in NMU-mediated glucose sensing and taste modulation remains an important question for future investigation.

The conservation of this “satiety brake” suggests that suppressing sensory gain via central neuropeptidergic energy sensors is a fundamental strategy for maintaining energy balance. By defining a direct link between internal glucose levels and sweet taste perception, our work provides a mechanistic framework for understanding how metabolic state shapes sensory experience and feeding behavior.

## Materials and methods

### Animals

Fly: Flies were maintained on a standard fly diet under conditions of 60% humidity and 25 °C with a 12-hour light and 12-hour dark cycle. Virgin female flies were chosen immediately after emerging and housed in standard fly food (ND) vials, with 20 flies per vial, for a period of 5-6 days before experiments. For temperature sensitive manipulations (dTrpA1, Shibire^ts^), flies were reared at 18 °C for 7–8 days, then transferred to 30 °C for 30 min to activate or inhibit targeted neurons; behavioral assays (PER or feeding) were conducted continuously at 30 °C. For starvation, the flies were kept on 2 % Agar for 12-hour. For refeeding, the fasted flies were transfer to different food for 15 mins.

The following UAS-RNAi lines: *UAS-AstA-R1 RNAi* (#27280), *UAS-AstA-R2 RNAi* (#25935), *UAS-PK2-R1 RNAi (#29624)*, *UAS-PK2-R2 RNAi (#28781)*, *UAS-CD8-GFP, LexAop-CD2-RFP (*#*67093)*, *UAS-GCaMP6m* (*#42748*), *UAS-CaMPARI* (*#58764*), *Gr5a-GAL4* (*#57592*), *Gr5a-LexA* (#93014), *ilp2-GAL4 (#37516)*, *elav-GAL4 (#25750)* were obtained from the Bloomington Drosophila Stock Center at Indiana University. *TH-KO*, *PK2-R1-KO, AstA-R1-KO* and all the other Gal4 and LexA lines (*AstA-Gal4, LK-Gal4, hugin-Gal4, TK-Gal4, CCHa1-Gal4, CCHa2-Gal4, TH-Gal4, PK2-R1-Gal4, AstA-LexA, AstA-R1-LexA*) were from Yi Rao (Deng et al., 2019) (Capital Medical University, Beijing). *UAS-dTrpA1* and *UAS-Shibire^ts1^* were from David Anderson (Caltech). *UAS-GCaMP6m; LexAop-CsChrimson* was from Yufeng Pan (Southeast University, Nanjing). *LexAop-Chrimson* was from Wei Zhang (Tsinghua University, Beijing).

Mouse: Male C57BL/6J mice at the age of four weeks were procured from cyagen. The mice were kept in standard laboratory conditions at a temperature of 25 with a 12:12 light/dark cycle. All experimental procedures were conducted in compliance with the guidelines set by the Laboratory Animal Welfare and Ethics Committee of Chongqing University (CQU-IACUC-RE-202401-004), adhering to both national and international standards.

### Behavioral assays (Fly)

PER was evaluated using a method outlined in a previous study{Marella, 2012 #22}. Briefly, individual flies were gently aspirated and positioned in a 200 µL pipette tip, allowing only the head and proboscis to protrude while the dorsal thorax was lightly affixed to prevent escape. After a 3-minute acclimation period, each fly was first presented with 0.5 µL water droplets on the labellum twice. Flies that responded by extending their proboscis were allowed to drink and were subsequently excluded from the experiment. Flies that did not extend their proboscis in response to water were then stimulated sequentially with ascending sucrose concentrations (6.25–800 mM). Each concentration was delivered as two brief (<1 s) touches of a 0.5 µL droplet to the labellum, immediately withdrawn to prevent ingestion. A full proboscis extension, defined as the complete uncoiling of the rostrum beyond the labellum plane, was scored as positive. Partial or delayed (>1 s) extensions were not considered positive responses. Responsiveness at each concentration was recorded if at least one full extension occurred in the two trials.For Figure supplementary 11, the connection between the brain and VNC were cut off by a dissecting scissors before the PER assay.

Feeding was assessed as previously described ^78^. Briefly, 5-day-old flies underwent a 12-hour starvation period on 2% agar and were then moved to a new vial with test food containing 0.5% Brilliant Blue (MACKLIN, China) for 10 minutes. After a rapid freeze at -20 °C, flies were decapitated in PBS, homogenized, and centrifuged (13,000 g * 5 min). The resulting supernatants were diluted with PBS to a total volume of 1 ml, and absorbance was measured at 620 nm.

### Sucrose Preference Test

The two-bottle preference (TBP) behavioral test was conducted in a standard mouse cage. Two bottles, one containing water and the other with a 10 mM sucrose solution, were placed on top of the cage. Each tested mouse was individually caged and adopted to two bottles containing distilled water or a 10 mM sucrose solution for one week. Throughout the training period, the positions of the bottles were alternated every 24 hours to mitigate potential position-related biases. Prior to the TBP tests for sucrose, mice underwent a 12-hour pre-starvation period, followed by re-fed sucrose (20%) for 1 hour or intraperitoneal (IP) administration of NMU (YFLFRPRN-NH_2_, 4.5 µm/kg). The consumption of water and sucrose solution over a 2-hour period was meticulously recorded. The sweetener preference ratio was then calculated by dividing the weight of sweetener consumed by the weight of water consumed.

### NMU measurement

Blood was obtained from the lower jaw of the mice and clotted at room temperature for 1 hour. The samples were then centrifuged (1,500g, 4 °C) for 10 minutes and the supernatants were collected for NMU measurement. All the manipulations were according to the manufacturer’s instructions (BMASSY,74030).

### Hemolymph extraction and glucose measurement

Groups of 40 ice anesthetized flies were decapitated and placed into a perforated 0.5 ml tube, which was nested inside a 1.5 ml collection tube. Samples were centrifuged at 2,500 × g for 10 min at 4 °C, yielding approximately 2 µl of hemolymph per batch. Hemolymph was pooled to 4 µl per assay (n = 8 independent samples per group) and immediately analyzed for glucose concentration using the Solarbio BC2500 kit (Solarbio, China) according to the manufacturer’s instructions.

### Microinjection

Flies were delicately positioned and secured within a 200 ml pipette tip, ensuring their heads were directed toward the tip’s end. Subsequently, the tip was opened by cutting, exposing the heads and a section of the thorax. About 20 nl of sterile saline, either with or without synthesized peptides (hugin: SVPFKPRL-NH_2_, 1 mg/ml (1.1 mM); AstA: LPVYNFGL-NH_2_, 1mg/ml (1.1 mM); NMU: YFLFRPRN-NH_2_, 1mg/ml (0.9 mM); Control peptide: DYKDDDDKYPYDVPDYA, 1mg/ml (0.48 mM), were injected into the thorax of these flies using a glass micropipette and a microinjector (3-000-207, Drummond Scientific Company Instruments). The glass micropipette was created from thick-walled borosilicate capillaries (3-000-203-G/X, Drummond Scientific Company Instruments).

### Immunofluorescence staining

Fly brains were carefully dissected in PBS on ice and then fixed in 4% formaldehyde for 60 minutes. Following fixation, the brains underwent permeabilization and blocking using Dilution/Blocking Buffer (PBS containing 10% Calf Serum and 0.5% Triton X-100) for 2 hours at room temperature. Subsequently, the samples were immersed in the appropriate primary antibodies in Dilution/Blocking Buffer for 24 hours at 4 °C. Afterward, the samples were subjected to a 60-minute wash with Washing Buffer (0.5% Triton X-100 in PBS) four times at room temperature and were then incubated with secondary antibodies for 24 hours at 4 °C. The samples underwent three additional washes with Washing Buffer before being mounted in Fluoroshield (Sigma-Aldrich).

Images were acquired using a scanning confocal microscope with Olympus objectives (20× /0.7 and 40× /0.95w). Antibodies were employed at the following dilutions: mouse anti-nc82 (1:100, DSHB), rabbit anti-GFP (1:500, Abcam), Alexa Fluor 647 goat anti-mouse (1:500, Invitrogen), and Alexa Fluor 488 goat anti-rabbit (1:500, Invitrogen).

### In situ hybridization

Mice were anesthetized and perfused with 4% paraformaldehyde (PFA) dissolved in phosphate-buffered saline (PBS, pH 7.4). Brains were carefully removed from the skull and post-fixed in 4% PFA at 4 °C overnight. Subsequently, the brains were transferred to a 20% sucrose solution (in 1 × PBS) at 4 ° C until they were ready for sectioning. Twenty-micrometer-thick sections were obtained using a cryostat. The detection of nmu was performed using a fluorescent in situ hybridization technique (RNAscope, Pinpoease) according to the manufacturer’s instructions. Sections were mounted on SuperFrost Plus Gold slides (ThermoFisher) and briefly rinsed in autoclaved Millipore water. They were then subjected to gradient dehydration in 50%, 75%, and 100% ethanol for 5 minutes each. A hydrophobic barrier was created around the sections using an ImmEdge hydrophobic barrier pen (Cat No. 310018). All incubation steps were carried out at 40 °C using the Pinpoease hybridization system (Cat No. SH-08). The subsequent hybridization, amplification, and detection steps were performed strictly according to the manufacturer’s instructions.

### Stereotaxic brain surgeries and viral injection

Mice were anesthetized with 1-2% isoflurane during stereotaxic injections, performed using a small animal stereotaxic instrument (RWD Life Science, #68030). Throughout the surgery, core body temperature was maintained at 36 ± 1 °C using a feedback-controlled heating system. Micro scissors were used for the incision, and a dental drill was employed to create the cranial window. Viral injections were delivered using a glass microelectrode syringe pump (RWD Life Science, #R-480). To express GCaMP6m and mCherry AAV2/9-EF1α-DIO-GCaMP6m-WPRE (titer: 2.89 × 10^12^, #H4955, Obio Biotechnology, Shanghai, China, Co., Ltd.) and AAV2/9-EF1α-DIO-mCherry-WPRE (titer: 2.89 × 10^12^, Obio Biotechnology, Shanghai, China, Co., Ltd.) were bilaterally injected into the VMH and rNST, with approximately 0.15 µL of virus delivered per site. The microelectrode was left in place for 10 minutes post-injection before being withdrawn. Injection coordinates were as follows: VMH (AP, −1.46 mm; ML, ±0.4 mm; DV, −5.5 mm) and rNST (AP, −7.08 mm; ML, ±0.6 mm; DV, −4.3 mm). After 3 weeks of AAV injections, optical fibers were chronically implanted in the VMH or rNST and secured with dental cement (C&B Metabond®, Parkell, Japan). Injection coordinates were determined based on the 4th edition of the mouse brain atlas by Franklin and Paxinos (2008). Mice were allowed to recover for 5-7 days post fiber-implanted before any behavioral or calcium imaging evaluations were conducted. After the experiments, mice were perfused to verify virus expression and fiber placement. Data from mice with poor viral expression or incorrect fiber placement (0-20% of cases) were excluded from the final analysis.

For the anterograde tracing experiments, two viral vectors were used. AAV2/1-hsyn-DIO-EGFP-WPRE (titer: 1 × 10^13^ viral particles/mL,Obio Biotechnology) was injected into the ventromedial hypothalamus (VMH), and AAV2/1-hsyn-SV40 NLS-Cre (titer: 2.1 × 10^12^ viral particles/mL, Brain Case) was injected into the rostral nucleus of the solitary tract (rNST). Approximately 0.15 µL of virus was delivered per injection site. After 3 weeks, the brains were perfused and fixed. Subsequently, the brain tissues were sliced and subjected to antibody staining. The fluorescence signals were then recorded and analyzed.

### Optical-fiber-based calcium recording in freely mice

Following injection of an AAV2/9-EF1α-DIO-GCaMP6m-WPRE viral vector, an optical fiber (200 μm O.D., 0.37 numerical aperture, Inper Inc., China) was placed 150 μm above the viral injection site. GCaMP6m^+^ mice were implanted with the optic fiber 3 weeks post-AAV injection and allowed to recover for 5–7 days prior to experimental testing.

For Figure 5K-L, six mice were first trained to lick glucose (500 mM) from a petri dish. After training, they were fasted for 12 hours and then received an intraperitoneal injection (ip) of synthetic NMU peptide (4.5 µm/kg), followed by a 30-minute waiting period. During the experimental session, the mice were given 10 seconds to lick glucose from the petri dish while undergoing fiber photometry recording. For Figure 5E-G, mice were fasted for 4 hours and then administered glucose via gastric infusion, with fiber photometry recordings capturing the process. Fluorescence signals were acquired using a dual-channel fiber photometry system (410 nm & 470 nm, RWD Life Science). The laser power at the tip of the optical fiber was kept below 20 μW for both the 470 nm and 410 nm channels to minimize bleaching. ΔF / F was calculated as (470 nm signal - fitted 410 nm signal) / fitted 410 nm signal. Data analysis was performed using software from RWD Life Science.

### Ex vivo calcium imaging

For Figure 1G, 1I, figure supplementary 6, and Figure 2 freshly dissected fly brains were placed in the sugar-free AHL buffer (108 mM NaCl, 8.2 mM MgCl_2_, 4 mM NaHCO_3_, 1 mM NaH_2_PO_4_, 5 mM KCl, 2 mM CaCl_2_, 5 mM HEPES, pH 7.3). Baseline recordings of the samples in AHL buffer were taken over 1 minutes. Subsequently, the solutions were added with 100 μM glibenclamide (***Figure 2D***), 50 mM pyruvate (***Figure 2G***) or switched to AHL + glucose (50 mM) with or without indicated chemicals (***Figure 2 A and E***, 1 mM phlorizin or 10 um alloxan), and the pH was adjusted back to 7.3 through gentle perfusion for 3 minutes. Solution flow in the perfusion chamber was controlled by a valve commander (Scientific Instruments). Following stimulation, samples were washed out again with AHL. For the TTX test, 2 μM TTX was added to the AHL solution. Prior to the assay, the samples were pre-treated with 2 mM TTX for 15 minutes.

For Figure 7H-J, mice (6-9 weeks of age) were deeply anesthetized by avertin (1.25% avertin, 0.2 ml/10g). Mouse brains were quickly extracted following decapitation and immediately placed in ice-cold slicing buffer (110 mM Choline Cl, 2.5 mM KCl, 1 mM NaH_2_PO_4_, 25 mM NaHCO_3_, 5 mM glucose, 7 mM MgCl_2_, 0.5 mM CaCl_2_,1.3mM Na ascorbate,0.6mM Na pyruvate bubbled with 95% oxygen and 5% CO_2_ with an adjusted pH of 7.3). The brains were then sectioned into 200 μm slices using a vibratome (Leica VT1200S). These slices were incubated artificial cerebrospinal fluid (aCSF) (125 mM NaCl, 2.5 mM KCl, 1 mM NaH_2_PO_4_, 25 mM NaHCO_3_, 5 mM glucose, 1.3 mM MgCl_2_, 2 mM CaCl_2_,1.3mM Na ascorbate, 0.6 mM Na pyruvate bubbled with 95% oxygen and 5% CO_2_ with an adjusted pH of 7.3) at 34 °C for 20 minutes, then transferred to a room temperature for 30 minutes before recording. For Ca^2+^ imaging, the slices were placed in a recording chamber in the low sugar aCSF (125 mM NaCl, 2.5 mM KCl, 1 mM NaH_2_PO_4_, 25 mM NaHCO_3_, 1 mM glucose, 1.3 mM MgCl_2_, 2 mM CaCl_2_, 1.3 mM Na ascorbate, 0.6mM Na pyruvate, pH 7.3) for 10 minutes (maintained the flow rate for 1-3 ml/ minute). Baseline recordings, perfusion, and Ca^2+^ recording were performed as detailed above.

All imaging was conducted on an Olympus confocal microscope (FVMPE-RS) with a water immersion objective lens (25× /1.05w MP). Image analyses were performed using ImageJ to calculate the mean intensity of the indicated neuron targeting ROIs, and then plotted in Excel (Microsoft). Ratio changes were calculated using the formula: ΔF / F = [F – F_0_] / F_0_, where F represents the mean fluorescence of the cell body, and F_0_ is the average baseline (before stimulation).

### Optogenetics and in vivo calcium imaging

Newly emerged virgin female flies were housed in a fresh vial with standard medium for 3 days and subsequently moved to a vial containing regular food supplemented with 400 µM all-trans-retinal (Sigma R2500) for 2-4 days prior to experiments. These flies were then immobilized on ice, affixed to transparent tape, and the dorsal cuticle of the fly head was delicately removed with forceps to expose the brain, which was immersed in AHL.

For Figure 3F, an array of red LED (Thorlabs, M625L3) was positioned above the brain. Calcium signals of Gr5a^+^ neurons were captured for 60 frames (1 second per frame) in the absence of light to establish a baseline. Subsequently, the next 20 frames were recorded during opto-activation with the light (the light was switched on for 0.45 second and rested for 0.55 second per activation cycle, and the images were collected in rest time), followed by another 60 frames recorded without light afterward.

For Figure 4G and 4H, to apply liquid food (5% sucrose) to the proboscis, a pipette was filled with the taste solution and placed a few microns from the proboscis tip by using a micromanipulator (MP225, Sutter Instruments) prior to recording. Then, we recorded calcium signals for 60 seconds in the absence of light to establish a baseline (1 second per frame). To activate neurons expressing CsChrimson, a red LED was positioned above the brain, controlled by a custom computer program during opto-activation (0.45 second on and 0.55 second off from 60 second to 120 second). The pipette was placed on the proboscis from 80 second to 85 second. Calcium signals of Gr5a^+^ neurons were recorded throughout these processes (1 second per frame).

Image analyses about relative fluorescence changes ΔF / F were performed as ex vivo calcium imaging.

### Optogenetics and ex vivo calcium imaging

For Figure supplementary 11C, newly emerged virgin female flies were housed in a fresh vial with standard medium for 3 days and subsequently moved to a vial containing regular food supplemented with 400 µM all-trans-retinal (Sigma R2500) for 2-4 days prior to experiments. The brains were dissected and placed under a red LED. Calcium signals of AstA^+^ neurons were recorded with or without LED light as above. Image analyses were performed as ex vivo calcium imaging.

### Calcium imaging with CaMPARI

The flies were then gently immobilized on ice, attached to transparent tape, and the dorsal cuticle of the fly head was carefully removed with forceps to expose the brain, which was immersed in AHL. Photoconversion (PC) was achieved for 30 seconds using a 405 nm LED [Thorlabs, M405L2–UV (405 nm) Mounted LED, 1,000 mA, 410 mW], controlled by a LED controller (Thorlabs, LEDDB1 driven with 1000 mA). Images were captured using an Olympus confocal microscope. Image analyses were performed in ImageJ by manually drawing ROIs covering individual neuronal cell bodies using the green channel. The extent of CaMPARI photoconversion was determined as the Red: Green ratio.A ‘-405 nm’ control was included to demonstrate that scanning of the hugin neurons without exposure to ultraviolet light does not convert green to red fluorescence.

### Statistical analysis

Data are shown as means (± SEM). Data presented in this study were verified for normal distribution by D’Agostino–Pearson omnibus test. Student’s t test, one-way ANOVA and two-way ANOVA (for comparisons among three or more groups and comparisons with more than one variant) were used. The post hoc test with Bonferroni correction was performed for multiple comparisons following ANOVA.

## Acknowledgements

We express gratitude to all members of the Wang and Huang Lab for their valuable discussions and technical support. Special thanks to Yi Rao and Bowen Deng for generously providing *Drosophila* chemoconnectome lines, and to Yulong Li and Jianzhi Zeng for their assistance with *in vivo* calcium imaging preparations. Appreciation is extended to the members of the Neuroscience Pioneer Club for their insightful discussions throughout the course of this study. This study was funded by the National Natural Science Foundation of China (No. 32071006 for L.W. and No. 32271008 for R.H.), Guangdong Basic and Applied Basic Research Foundation (2023A1515110454 for T.S.), and the startup funds from Shenzhen Bay Laboratory and Chinese Institutes for Medical Research to L.W.

## Author contributions

T.S., R.H., and L.W. designed the experiments and wrote the manuscript. W.Q., and T.S. conducted and analyzed all the experiments. L.W. and R.H. supervised the project.

---

**Figure supplementary 1.**
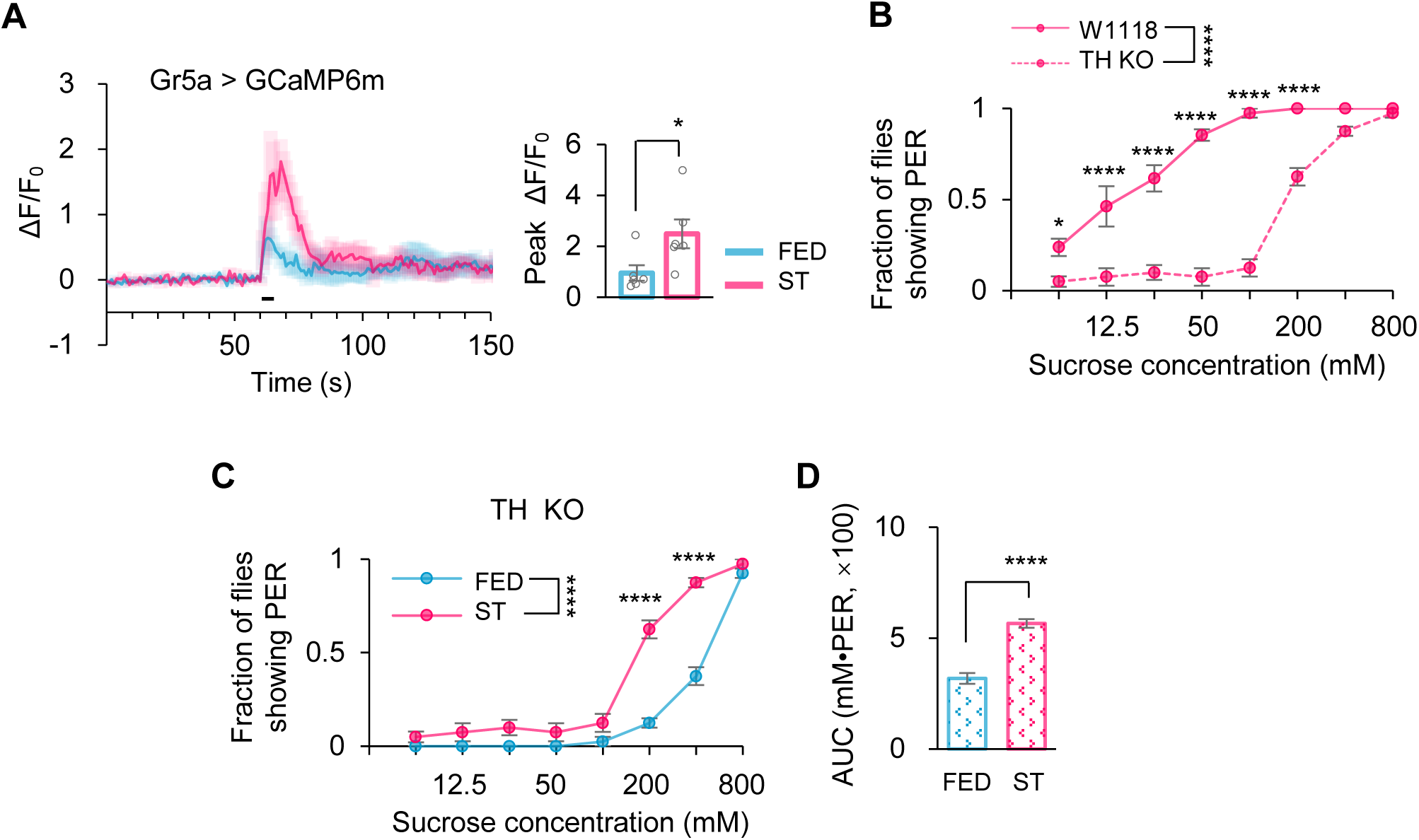
Satiety suppressed sweet sensitivity in a dopamine-independent manner. (A) Representative traces (left) and quantification (right) of peak calcium transients of Gr5a^+^ neurons in indicated flies (t-test; *, p=0.0373; n=6). Horizontal black bar represents the duration 5% sucrose stimulation. (**B-D**) Fraction of flies of the indicated genotypes showing PER to different concentrations of sucrose (two-way ANOVA; *, p=0.0498;****, p<0.0001; B-C, n=4 groups, each of 10 flies). The Area Under the Curve (AUC, D) represents sweet sensitivity in C. *P < 0.05; ****P < 0.0001. Student’s *t-test* and two-way ANOVA followed by post hoc test with Bonferroni correction were used for multiple comparisons when applicable.

**Figure supplementary 2.**
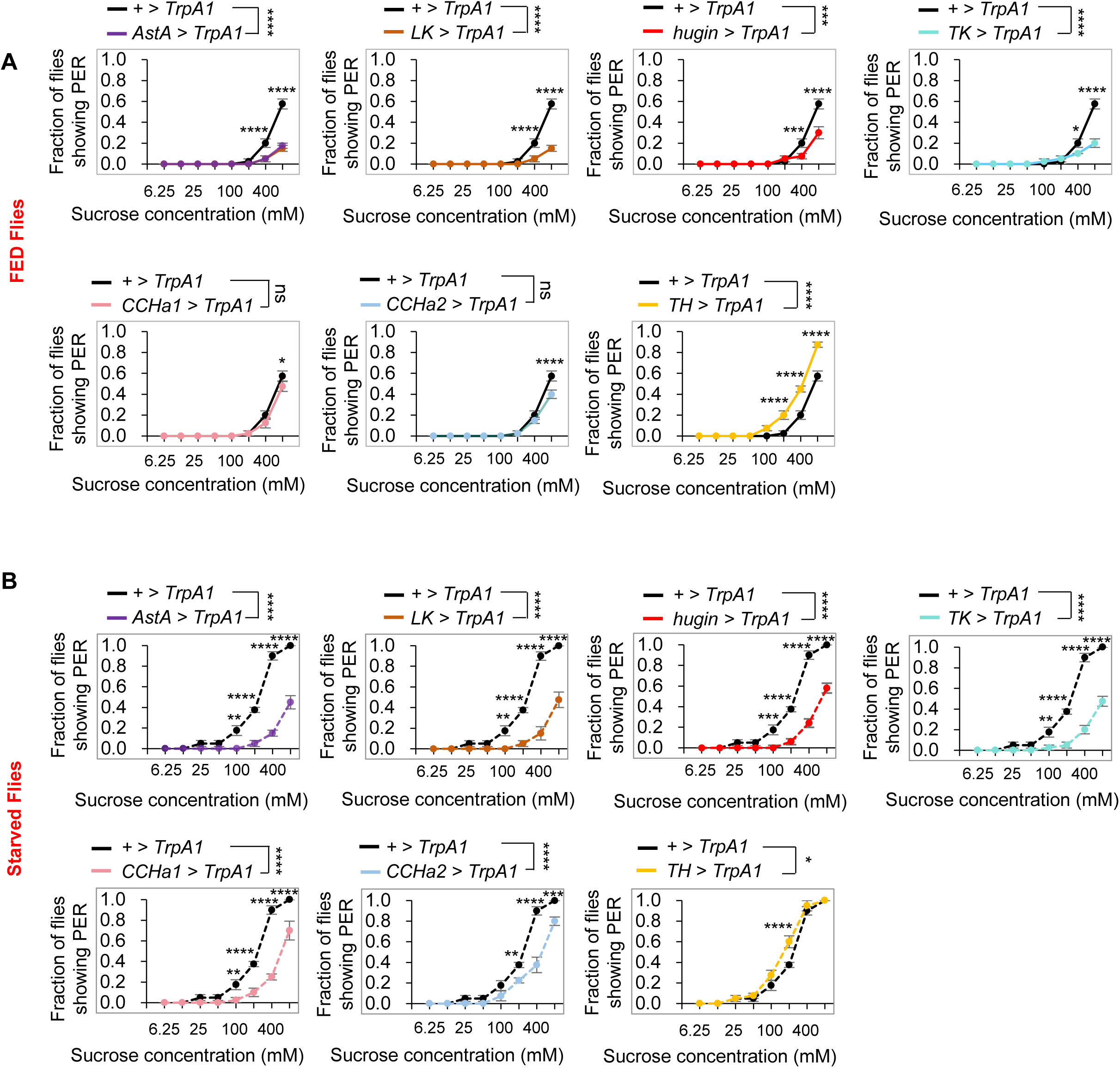
The effect of satiety signals on taste modulation. (A) Fraction of fed flies of the indicated genotypes and environmental temperatures showing PER to different concentrations of sucrose (two-way ANOVA; *, p=0.012; ***, p=0.0006;****, p<0.0001; n=4 groups, each of 10 flies). (**B**) Fraction of starved flies of the indicated genotypes and environmental temperatures showing PER to different concentrations of sucrose (two-way ANOVA; *, p=0.0154; **, p=0.0081 or 0.0011; ***,p=0.0005 or 0.0001; ****, p<0.0001; n=4-5 groups, each of 10 flies). ns, P > 0.05; *P < 0.05; ***P < 0.001; ****P < 0.0001. Two-way ANOVA followed by post hoc test with Bonferroni correction was used for multiple comparisons when applicable.

**Figure supplementary 3.**
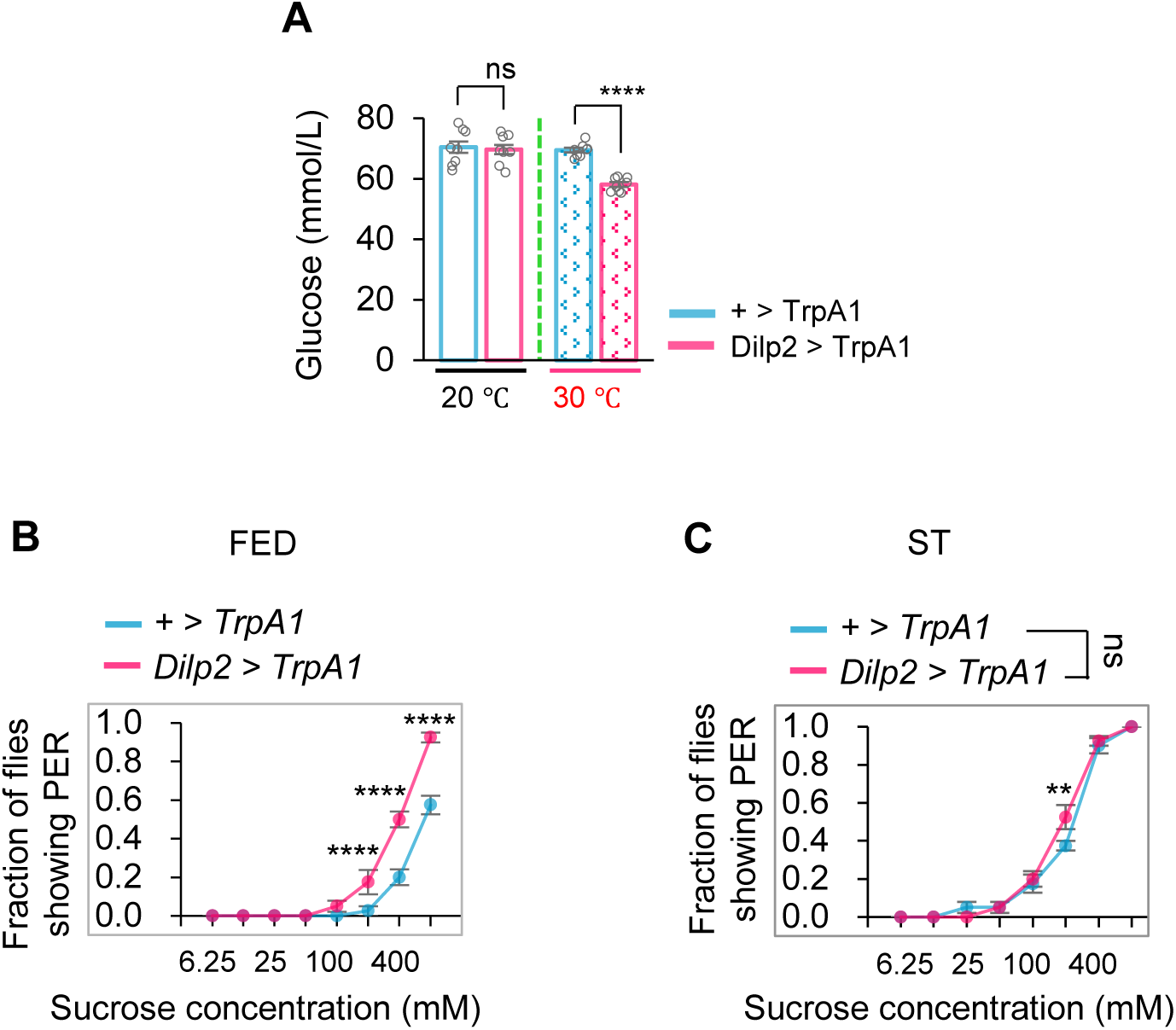
Activation of IPCs decreased the circulating sugar levels and increased sweet sensitivity. (A) Hemolymph glucose levels from the indicated genotypes and treatment temperatures (t-test; ****, p<0.0001; n=8-9, 1 hour at 30 °C). (**B-C**) Fraction of fed flies of the indicated genotypes and environmental temperatures showing PER to different concentrations of sucrose (two-way ANOVA;**, p=0.0048; ****, p<0.0001; n=4 groups, each of 10 flies). ns, P > 0.05; ****P < 0.0001. Student’s *t-test* and two-way ANOVA followed by post hoc test with Bonferroni correction were used for multiple comparisons when applicable.

**Figure supplementary 4.**
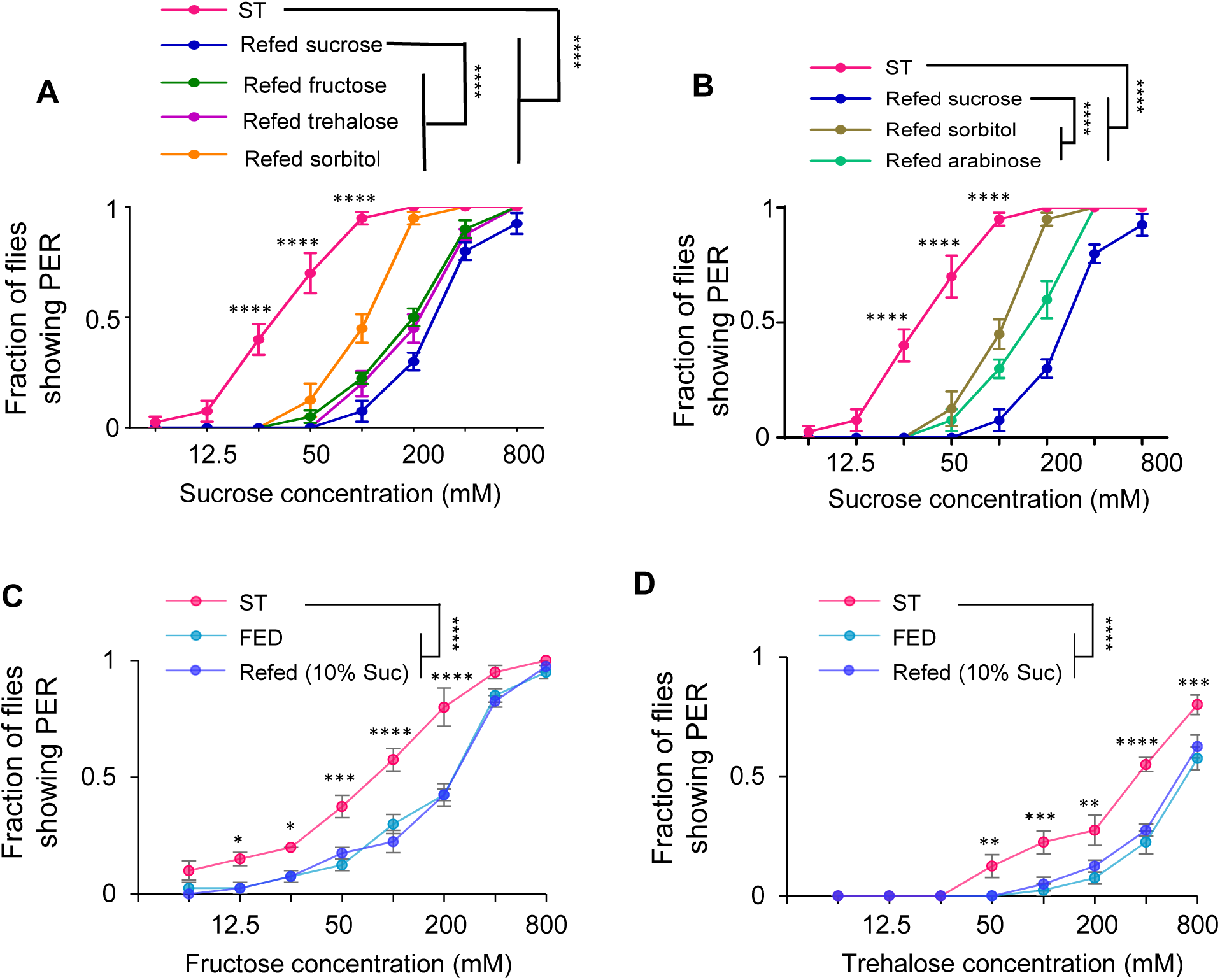
Sugar intake suppressed sweet sensation. (A) Fraction of indicated flies which refed with different sugar with energy showing PER to different concentrations of sucrose (two-way ANOVA; ****, p<0.0001; n=4 groups, each of 10 flies). (B) Fraction of indicated flies which refed with different sugar without energy showing PER to different concentrations of sucrose (two-way ANOVA; ****, p<0.0001; n=4 groups, each of 10 flies). (**C**) Fraction of indicated flies showing PER to different concentrations of fructose (two-way ANOVA; *, p=0.0255; ***, p=0.0002; ****, p<0.0001; n=4 groups, each of 10 flies). (**D**) Fraction of indicated flies showing PER to different concentrations of trehalose (two-way ANOVA;**, p=0.0081 or 0.0013; ***, p=0.0002; ****, p<0.0001; n=4 groups, each of 10 flies). ****P < 0.0001. two-way ANOVA followed by post hoc test with Bonferroni correction were used for multiple comparisons when applicable.

**Figure supplementary 5.**
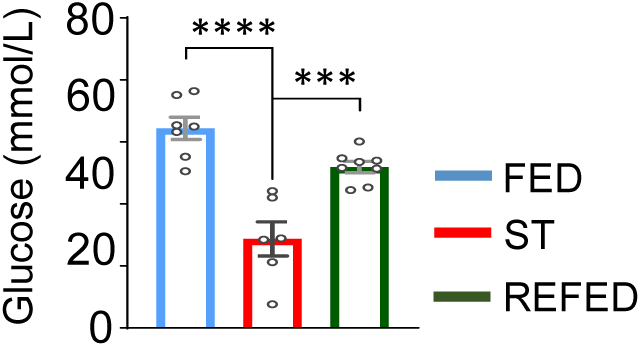
Starvation led to a decrease in the circulating sugar levels. Hemolymph glucose levels from indicated flies (t-test; ***, p=0.00017; ****, p<0.0001; n=6-8). ***P < 0.001. Student’s *t-test* was used for comparisons.

**Figure supplementary 6.**
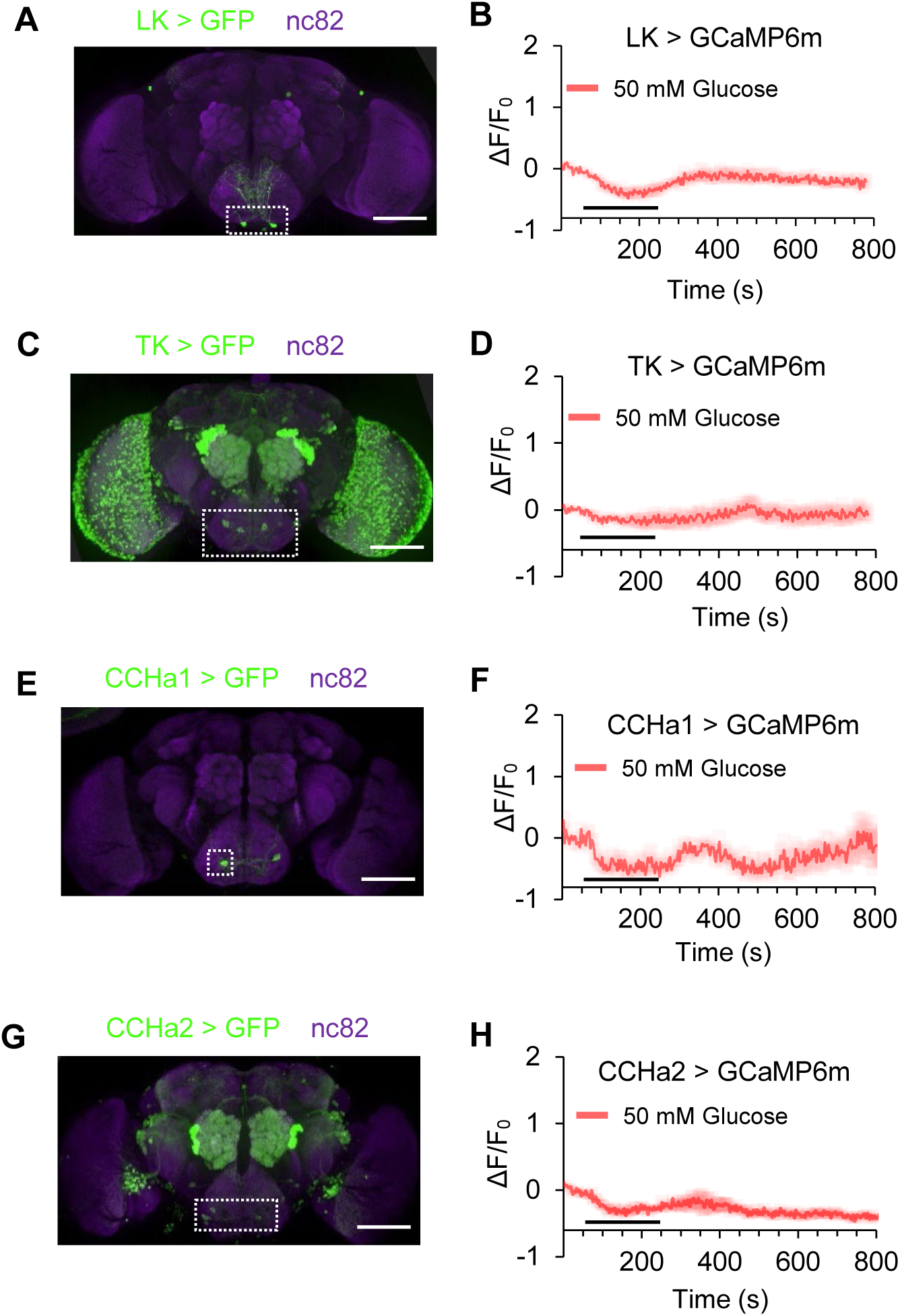
The response of different neuropeptide-expressing neurons to glucose. (**A**) LK expression in the brain, illustrated by mCD8::GFP expression driven by *LK^GAL4^*. Scale bar represents 100 μm. (**B**) Representative traces and quantification of ex vivo calcium responses of LK^+^ neurons during the perfusion of glucose (n=6). (**C-D**) The distribution (C) and calcium responses (D) of TK^+^ neurons (n=6). (**E-F**) The distribution (E) and calcium responses (F) of CCHa1^+^ neurons (n=6). (**G-H**) The distribution (G) and calcium responses (H) of CCHa2^+^ neurons (n=6).

**Figure supplementary 7.**
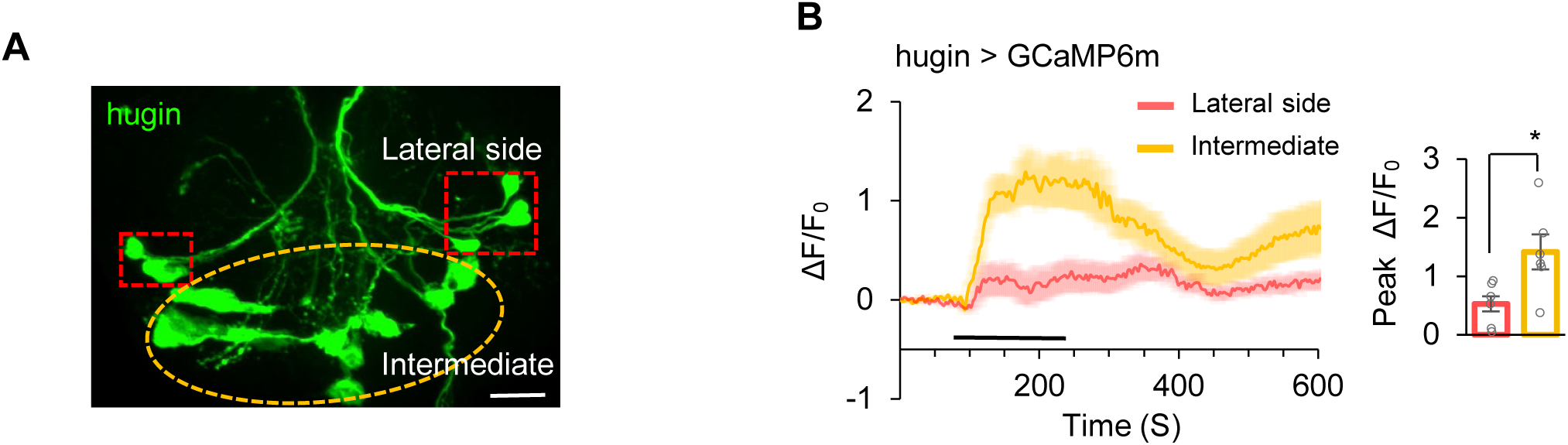
The calcium response of hugin neurons upon glucose stimuli. (**A**) Representative image of hugin neurons in SEZ. Scale bar represents 50 μm. (**B**) Calcium responses of different cluster of hugin neurons during the prefusion of glucose (t-test, *, p=0.0151; n=6).*P < 0.05; ***P < 0.001. Student’s *t-test* used for comparisons.

**Figure supplementary 8.**
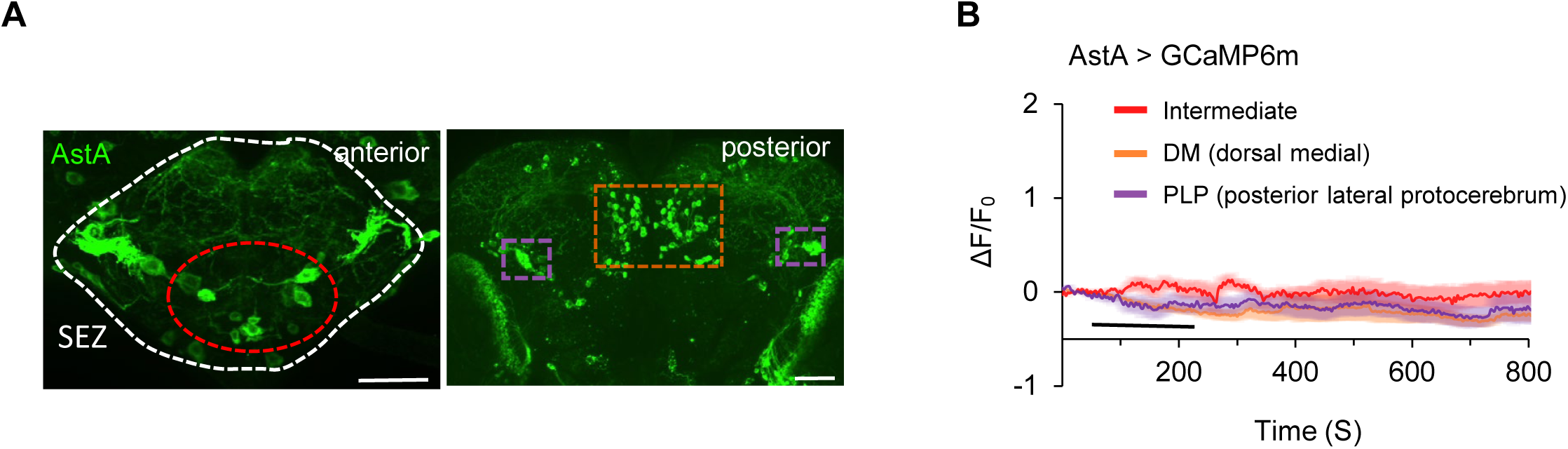
The calcium response of AstA^+^ neurons upon glucose treatment. (**A**) Representative image of different part of AstA neurons. Scale bar represents 50 μm. (**B**) Calcium responses of different cluster of hugin neurons during the prefusion of glucose (n=6)

**Figure supplementary 9.**
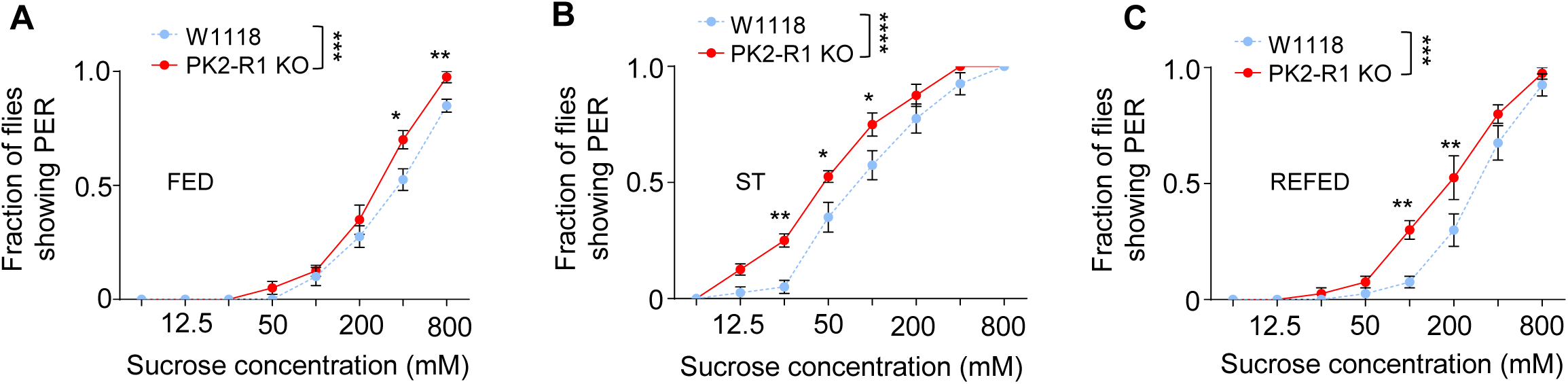
Knockout of *PK2-R1* enhanced sweet sensation. (**A-C**) Fraction of flies of the indicated genotypes showing PER to different concentrations of sucrose (two-way ANOVA; *, p=0.0464 or 0.0153; **, p=0.0015 or 0.0038 or 0.0035; ***, p= 0.0002; ****, p<0.0001; n=4 groups, each of 10 flies). ***P < 0.001; ****P < 0.0001. Two-way ANOVA followed by post hoc test with Bonferroni correction were used for multiple comparisons when applicable.

**Figure supplementary 10.**
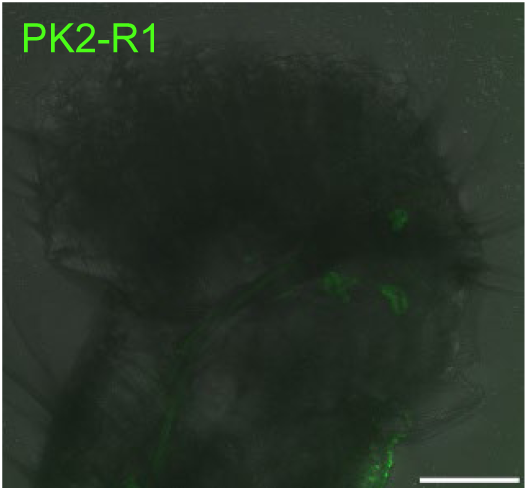
The expressing pattern of PK2-R1 in the proboscis. Representative image of PK2-R1 in the proboscis. Scale bar represents 50 μm.

**Figure supplementary 11.**
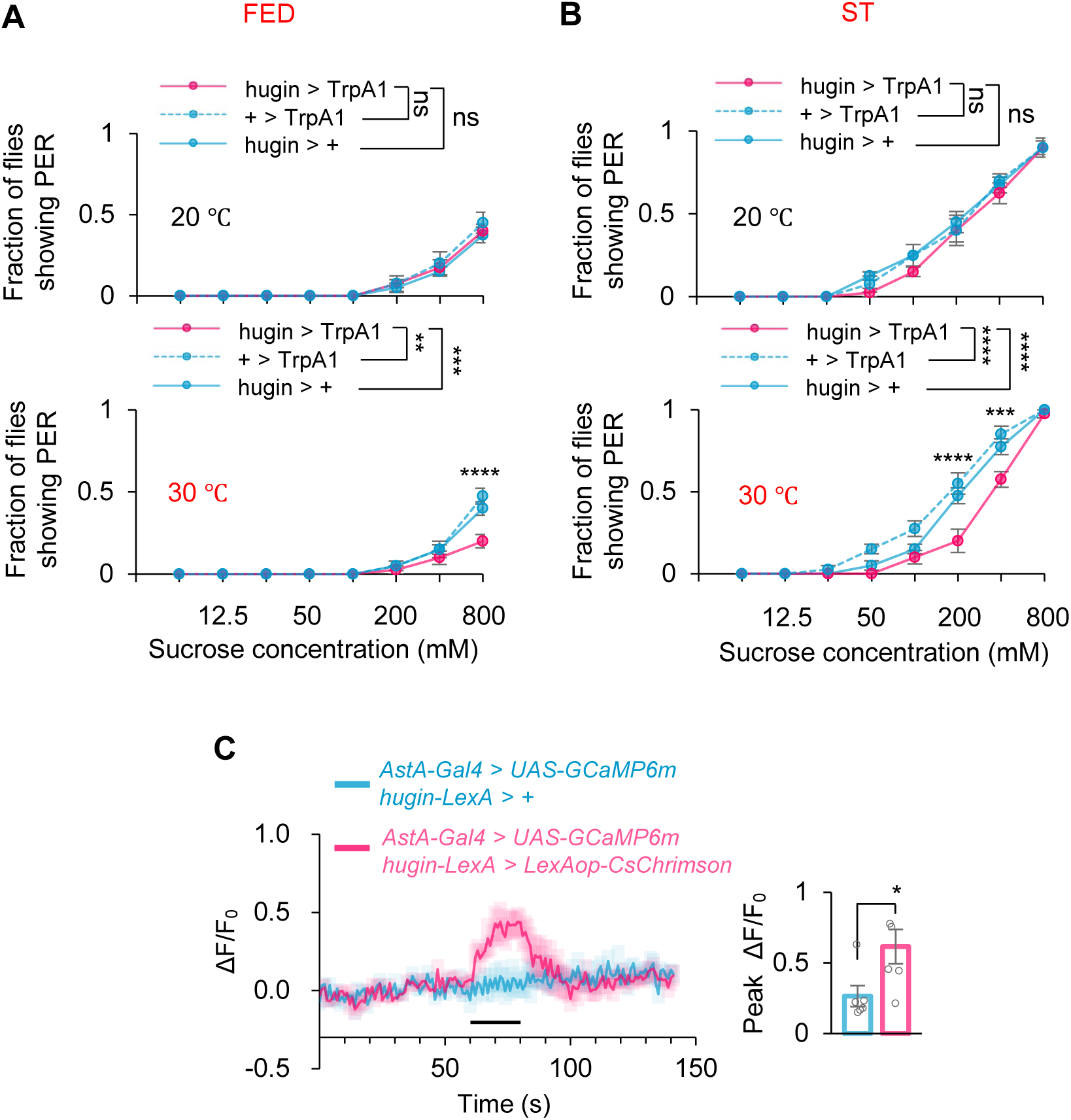
hugin^+^ neurons in the brain activated AstA^+^ neurons. (**A-B**) Fraction of flies of the indicated genotypes and environmental temperatures showing PER to different concentrations of sucrose (two-way ANOVA; **, p=0.0082; ***, p= 0.0007 or 0.0001; ****, p<0.0001; n=4 groups, each of 10 flies). Prior to the behavioral assay, the neural connection between the brain and the ventral nerve cord (VNC) was surgically severed, while the flies remained physically intact and retained normal proboscis musculature and motor output. PER was measured using the same protocol as in intact flies. (**C**) Representative traces (upper) and quantification (lower) of peak calcium transients of AstA^+^ neurons after the photo-activation of hugin^+^ neurons (t-test;*, p=0.0337; n=6) from ex vivo calcium imaging. Horizontal black bar represents the duration of red-light stimulation.

**Figure supplementary 12.**
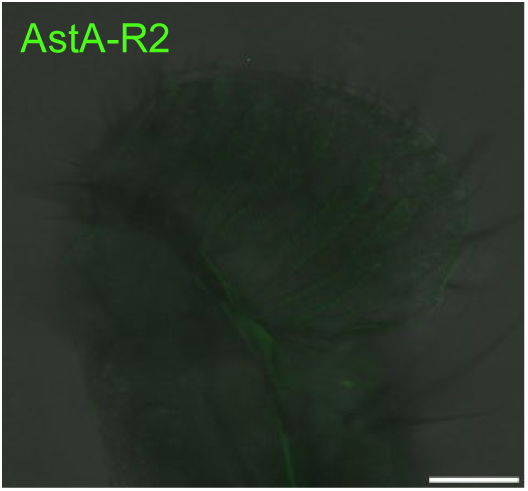
Manipulating gene *AstA-R1* enhance PER. (**A-C**) Fraction of flies of the indicated genotypes showing PER to different concentrations of sucrose (two-way ANOVA; *, p=0.0404; **, p=0.009 or 0.0022 or 0.0019;***, p=0.0005; ****, p < 0.0001; n=4 groups, each of 10 flies). ***P < 0.001; ****P < 0.0001. Two-way ANOVA followed by post hoc test with Bonferroni correction were used for multiple comparisons when applicable.

**Figure supplementary 13.**
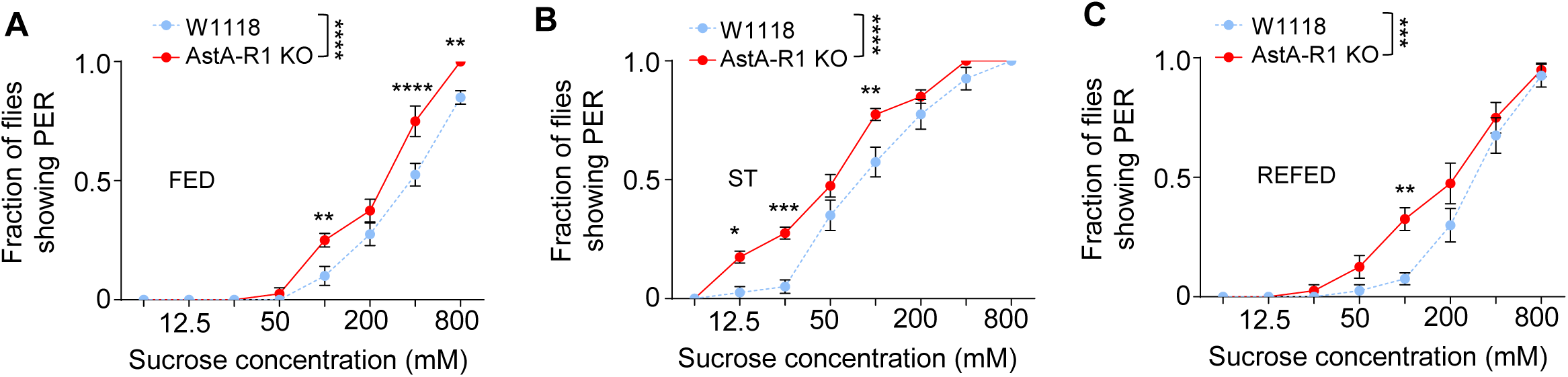
The expressing pattern of AstA-R2 in the proboscis. Representative image of AstA-R2 in the proboscis. Scale bar represents 50 μm.

**Figure supplementary 14.**
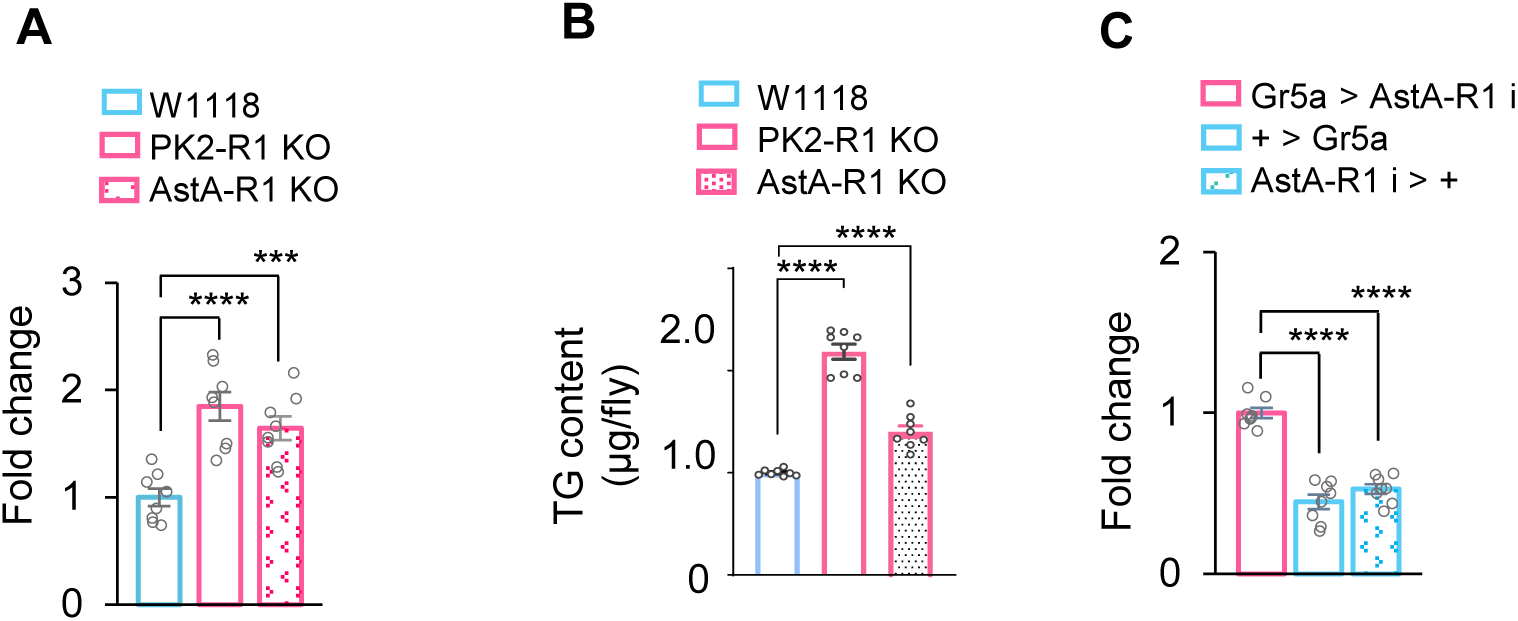
hugin-AstA-Gr5a circuitry inhibited feeding behavior. (**A**) Food consumption of flies of the indicated genotypes (one-way ANOVA; ***, p=0.0009; ****, p < 0.0001; n=8 groups, each of 4 flies). (**B**) The TG content of indicated genotypes (one-way ANOVA; ****,p<0.0001; n=8 groups, each of 2 flies). (**C**) Food consumption of flies of the indicated genotypes (one-way ANOVA; ****,p<0.0001; n=8 groups, each of 4 flies). ns, P > 0.05; ***P < 0.001; ****P < 0.0001. Student’s *t-test* and two-way ANOVA followed by post hoc test with Bonferroni correction were used for multiple comparisons when applicable.

**Figure supplementary 15.**
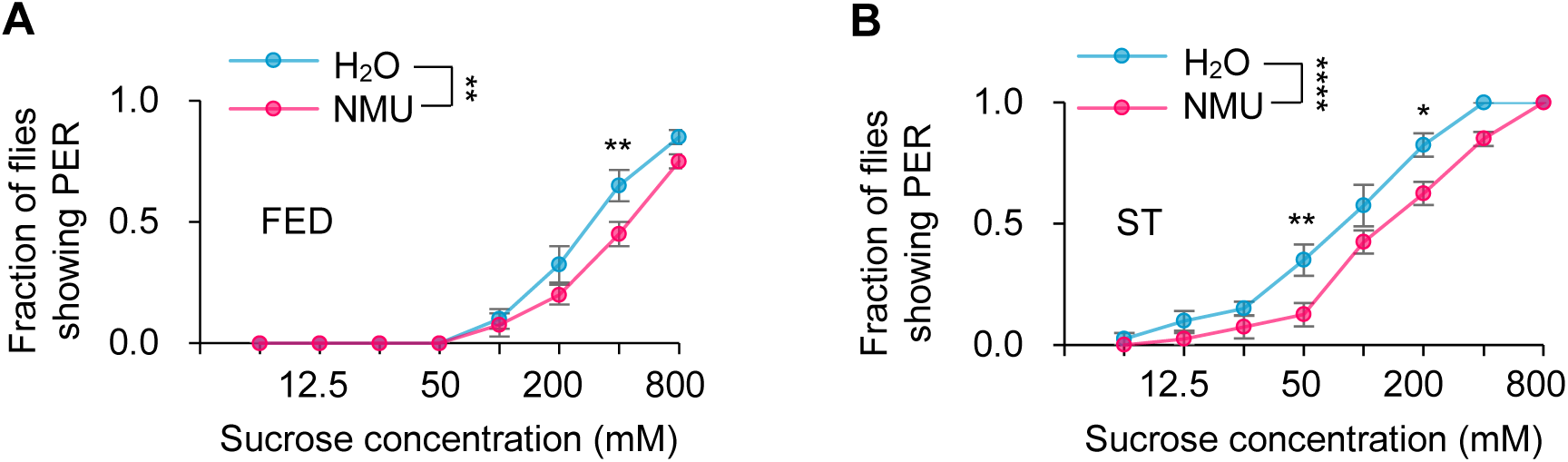
NMU peptide suppressed sweet taste in fly. Fraction of flies showing PER to different concentrations of sucrose (two-way ANOVA; *, p=0.106; **, p=0.0015 or 0.0029 or 0.0025; ****, p < 0.0001; n=4 groups, each of 10 flies). Flies were injected with saline or synthetic NMU for 30 mins before the assay.**P < 0.01; ****P < 0.0001. Two-way ANOVA followed by post hoc test with Bonferroni correction were used for multiple comparisons when applicable.

**Figure supplementary 16.**
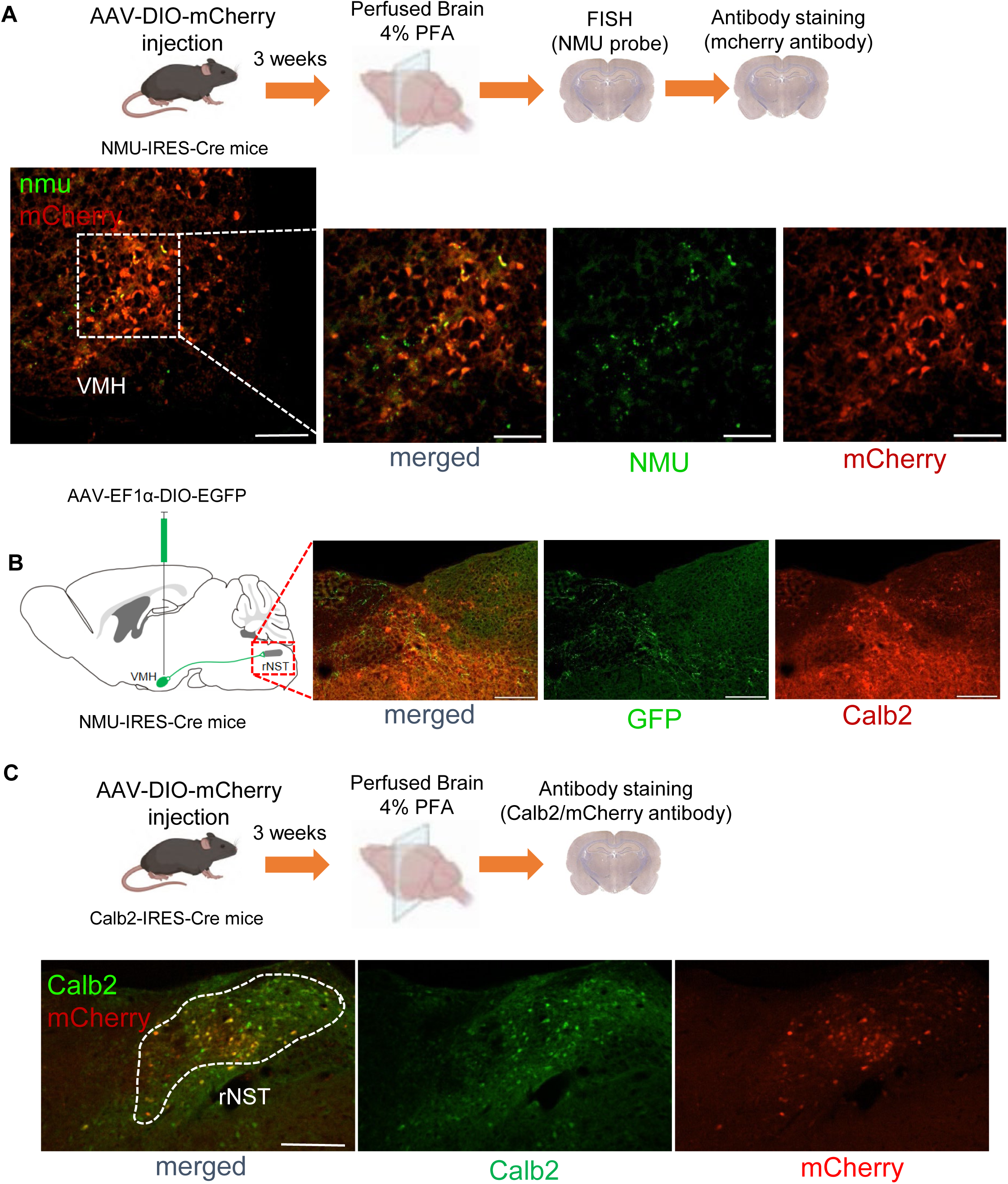
The expressing pattern of NMU neurons and Calb2 neurons. (**A**) Co-labeling of NMU^+^ neurons using a Cre-dependent mCherry reporter (red) and in situ hybridization (green).Scale bar represents 200 μm. (**B**) NMU arbors was extended near Calb2^rNST^ neurons. Scale bar represents 200 μm. (**C**) Co-labeling of Calb2^+^ neurons with a Cre-dependent mCherry reporter (red) and NMU antiboy staining (green). Scale bar represents 200 μm.

**Figure supplementary 17.**
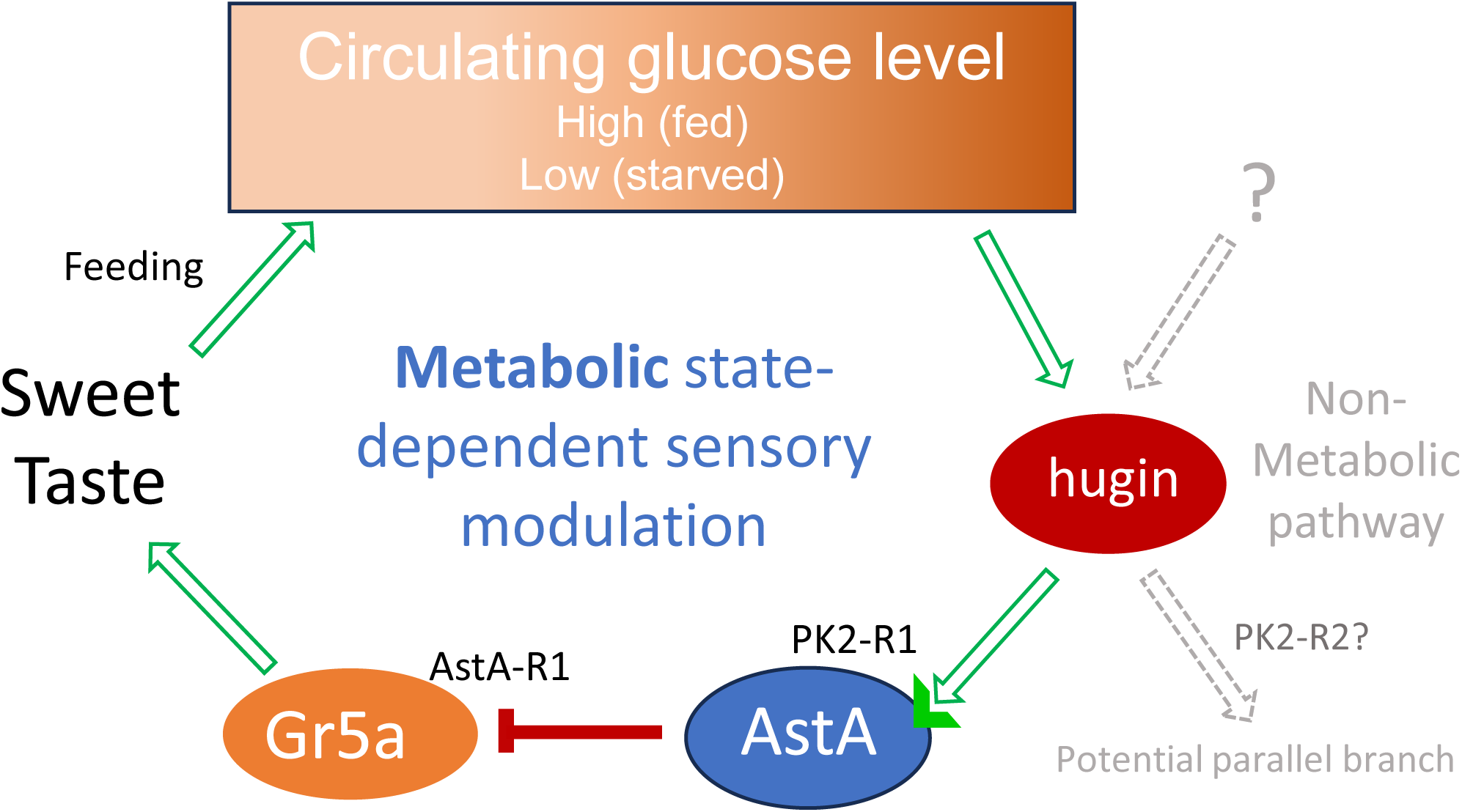
A working model. Elevated circulating glucose following feeding directly activates hugin+ neurons via cell-autonomous glucose sensing. Activated hugin neurons release Hugin neuropeptide, which engages PK2-R1 receptors on AstA neurons, promoting AstA release. AstA then suppresses the sensitivity of Gr5a-expressing gustatory sensory neurons through AstA-R1 signaling, thereby reducing sweet-driven feeding behavior. Under starvation, reduced circulating glucose decreases endogenous hugin neuronal activity, partially relieving this inhibitory tone and permitting enhanced sweet sensitivity. However, experimental manipulation of the circuit demonstrates that the hugin–AstA axis retains the capacity to modulate feeding behavior across feeding states, indicating that it functions as a glucose-responsive, state-modulated inhibitory system rather than a strictly satiety-exclusive brake. The modest phenotype observed upon PK2-R2 reduction suggests the potential existence of parallel or complementary hugin-dependent pathways, possibly arising from functional heterogeneity within hugin neuronal subpopulations. These additional branches remain to be defined.

